# Metabolic dynamics of the coral-algal symbiosis from fertilization to settlement identify critical coral energetic vulnerabilities

**DOI:** 10.1101/2023.03.20.533475

**Authors:** Ariana S. Huffmyer, Kevin H. Wong, Danielle M. Becker, Emma Strand, Tali Mass, Hollie M. Putnam

**Author notes:** Correspondence: Ariana S. Huffmyer.

## Abstract

Climate change accelerates coral reef decline and jeopardizes recruitment essential for ecosystem recovery. Adult corals rely on a vital nutritional exchange with their symbiotic algae (Symbiodiniaceae), but the dynamics and sensitivity of this reliance from fertilization to recruitment are understudied. We investigated the physiological, metabolomic, and transcriptomic changes across 13 developmental stages of *Montipora capitata*, a coral in Hawai‘i that inherits symbionts from parent to egg. We found that embryonic development depends on maternally provisioned mRNAs and lipids, with a rapid shift to symbiont-derived nutrition in swimming larvae. Symbiont density and photosynthesis peak once swimming to fuel pelagic larval dispersal. In contrast, respiratory demand increases significantly during metamorphosis, settlement, and calcification, reflecting this energy-intensive morphological reorganization. Symbiont ontogenetic proliferation is driven by symbiont ammonium assimilation with little evidence of nitrogen metabolism in the coral host. As development progresses, the host enhances nitrogen sequestration, regulating symbiont populations, and ensuring the transfer of fixed carbon to support metamorphosis, with both metabolomic and transcriptomic indicators of increased carbohydrate availability. Although algal symbiont communities remained stable, bacterial communities shifted with ontogeny, associated with holobiont metabolic reorganization. Our study reveals extensive metabolic changes during development, increasingly reliant on symbiont nutrition. Metamorphosis and settlement emerge as the most critical periods of energetic vulnerability to projected climate scenarios that destabilize symbiosis. This highly detailed elucidation of symbiotic nutritional exchange relative to sensitive early life stages provides essential knowledge for understanding and forecasting nutritional symbiosis integration, and specifically, coral survival and recruitment in a future of climate change.

## Introduction

Symbiotic relationships between animals and microbial organisms enable survival in nutritionally challenging environments ^1–4^. The function and structure of one of the most ecologically and economically important ecosystems on Earth, coral reefs, is driven by the productivity of marine symbioses. The nutritional symbiosis between reef-building corals and endosymbiotic algae in the family Symbiodiniaceae ^5^ can provide >100% of the coral’s daily metabolic demands through translocated organic carbon generated through symbiont photosynthesis ^6,7^. In return, metabolic waste from the coral host, including nitrogen and inorganic carbon, is available to Symbiodiniaceae to support their cellular metabolism and population growth ^8,9^. The energy-enhancing nutrient exchange between Symbiodiniaceae and corals, as well as their diverse bacterial and archaeal microbiome ^10^, enables coral biomineralization and thus the formation of the 3D skeletal structure that provides a habitat for high biodiversity in coral reef ecosystems ^11^ and their resulting invaluable goods and services for humans ^12^. However, the delicate coral-Symbiodiniaceae symbiosis is susceptible to destabilization by environmental stress. Rising sea surface temperatures and acute marine heat waves cause coral bleaching, or the dysbiosis and loss of Symbiodiniaceae cells and/or photosynthetic pigments, and reduced translocation of carbon surplus to the coral ^13^, which can, and often does, culminate in coral starvation and mass mortality ^14^. Increasingly, the stability of the nutritional relationship between corals, their algal endosymbionts, and symbiotic microbes is critical for the continued survival of coral reef ecosystems and the maintenance of essential ecosystem services humans rely on.

Globally, half of the live coral cover on reefs has been lost since the 1950s ^15^, significantly increasing the importance of coral reproduction and recruitment to replenish and maintain reef ecosystems. Corals have complex and biphasic life cycles, in which sessile adults produce gametes that are internally or externally fertilized and develop into swimming larvae that then undergo metamorphosis into the sessile polyp recruit stage and grow into mature colonies attached to the benthos ^16^. Survivorship from fertilization to recruitment represents the primary bottleneck for marine larvae ^17^, due to environmental impacts on fertilization, larval dispersal and behavior, predation, substrate suitability, and post-settlement survival ^16^. Key among the myriad challenges facing these sensitive life stages is the energetic requirement for successful development, metamorphosis, and calcification ^18^. To buffer the energetic challenges of living in oligotrophic waters, corals formed associations with Symbiodiniaceae ∼250 mya ^19^. This symbiotic nutritional exchange is initiated either immediately through vertical transmission (i.e., symbionts are provisioned to the eggs), or within days to weeks of release and/or settlement through the process of horizontal transmission (i.e., symbionts are acquired from the environment) ^20^. Initiation of symbiosis is a topic of intense study ^21^, but despite its increasing importance to reef persistence and recovery ^22^, the contribution of symbiont-derived nutrition to support energy demands of early life history are not well characterized. Broadly, an improved understanding of the development of symbiotic relationships in foundational marine invertebrates, like corals, provides critical insight on how environmental change may impact nutritional symbioses that are widespread across organisms and ecosystems.

Coral embryos and larvae initially metabolize limited endogenous lipid reserves ^23^, supporting the paradigm that parental provisioning is the primary energetic fuel for early life stages ^24–26^. For example, high-performance thin-layer chromatography lipid analysis in aposymbiotic life stages of corals (i.e., larvae) indicates a decline in wax ester abundance as larvae age and a strong reliance on parentally provisioned fatty acids for energetic fuel ^26^. Additionally, lipid usage in *Acropora tenuis* larvae without symbionts is highest within optimum thermal temperatures, suggesting that lipid stores are an important energy source for larval development and that dispersal potential will shift as ocean temperatures increase ^25^. However, in symbiotic larvae, the demand for energy fueled by lipid reserves has the potential to be substantially supplemented by photosynthetically derived carbon translocation. Radiolabel tracing in symbiotic *Pocillopora damicornis* larvae demonstrated that ∼70% of photosynthetically fixed carbon is translocated to the host ^27^. Further, the use of lipid reserves slows under light conditions in symbiotic *P. damicornis* and *Montipora digitata* larvae with lipids accounting for 38-41% of metabolic energy in the light compared to 84-90% of energy in the dark ^28^. Similarly, symbiotic *A. tenuis* larvae exhibited decreased lipid consumption compared to those that did not have symbionts, increasing dispersal potential and the ability of larvae to settle in favorable conditions ^29^. In contrast, high resolution local scale translocation using nanoscale secondary ion mass spectrometry (NanoSIMS) imaging combined with stable isotope in symbiotic *P. damicornis* larvae identified that the energetic contribution of photosynthates was systematically lower across tissues in larvae compared to the adult stage ^30^. These contrasting studies highlight the need to clarify the dynamics of nutritional exchange across early life history.

Recent evidence using stable isotope metabolomic tracing confirms that photosynthetically fixed carbon in the form of a major photosynthate, glucose, can be translocated to symbiotic larvae at approximately 7 days old (165 hours post fertilization) in *Montipora capitata* ^31^. Under thermal stress, *M. capitata* larvae increased nitrogen assimilation and sequestration to maintain glucose translocation, demonstrating that symbiotic relationships require metabolic investment to maintain ^31^. This previous research demonstrates that symbiotic relationships can contribute to nutritional requirements in coral larvae and have the potential to affect performance in coral larvae. However, it remains unclear at which point along development symbiotic nutritional exchange begins and the timing of transitions from reliance on lipids to active metabolism of photosynthates. Considering the need to increase our understanding of the sensitive coral symbiosis and energetic demands of recruitment, we examined physiological, molecular, and metabolomic data to characterize ontogenetic shifts in central metabolic pathways in early life stages of the vertically transmitting coral, *M. capitata* (Fig 1). Considering the energetic consequences of nutritional symbioses across taxa, here we examined an archetypal symbiotic nutritional model of reef-building corals to characterize: (1) symbiotic nutritional exchange relative to energetic demands across development, metamorphosis and settlement; and (2) mechanisms and dynamics of the transition from reliance on parentally provisioned lipid reserves to active utilization of symbiont nutrition.

**Fig 1.**
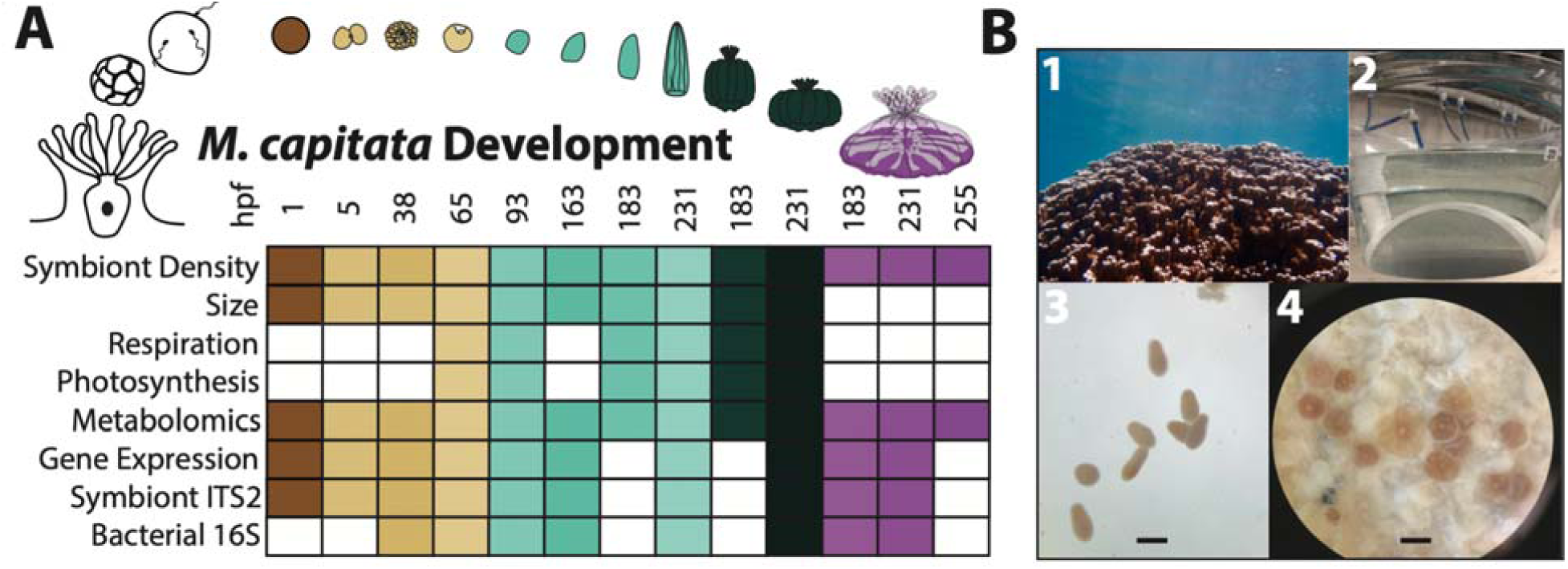
Characterization of *Montipora capitata* coral physiology and symbiosis across development. (A) Variables measured across *Montipora capitata* development. Filled squares indicate the response variable was sampled at the respective time point; white squares indicate the response was not measured. Color corresponds to major life history grouping (eggs=brown, embryos=yellow, larvae=cyan, metamorphosed polyps=green, attached recruits=pink). Hpf indicates hours post fertilization. (B) Photographs of 1 - *M. capitata* colony; 2 - Conical larval rearing chambers; 3 - Planula larvae, note pigmentation due to presence of symbiont cells, scale bar represents 1 mm; 4 - Attached recruits on settlement plug, scale bar represents 1 mm

## Results

### A. Ontogenetic shifts in physiology, metabolic rates, and endosymbiont photosynthesis

We tracked size, respiratory demand, symbiont density, and photosynthesis to characterize shifts in metabolic production and demand across development (65-231 hours post fertilization, hpf) (Fig 1AB). There were significant changes in individual size across development (Kruskal-Wallis, χ^2^(9,562)=235.86, P<0.001; Fig 2A). Embryo size decreased 41% as development progressed from egg (1 hpf) through larval stages as corals went through size contraction associated with the transition from the late gastrula (65 hpf) to planula (93 hpf) stages (Fig 2A). Size then increased, reaching a maximum at 231 hpf (142% growth relative to egg size; Fig 2A) in larvae and metamorphosed polyps.

**Fig 2.**
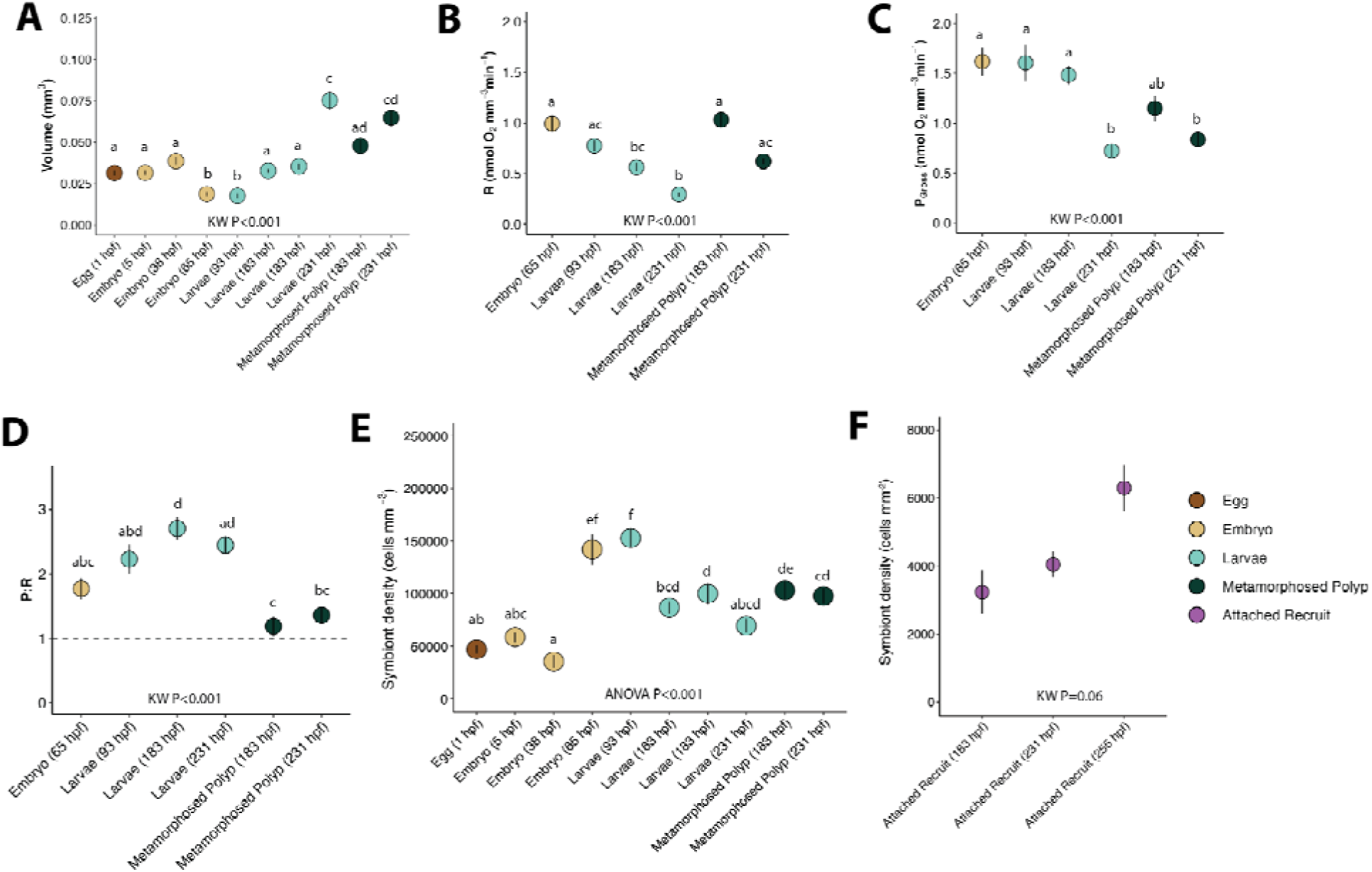
Metabolic rates and physiological responses change across *Montipora capitata* development. (A) Size (mm^3^) of each time point across development. (B) Size-normalized light enhanced dark respiration (R; oxygen consumed; µmol O_2_ mm^-3^ min^-1^). (C) Gross photosynthesis (oxygen produced; µmol O_2_ mm^-3^ min^-1^). (D) Gross photosynthesis : respiration ration (P:R). Dotted line indicates P:R of 1:1. (E) Symbiont cell density (cells per mm^3^) across development for egg through metamorphosed polyp stages. (F) Symbiont cell density (cells per mm^2^) in attached recruits. Data are represented as mean ± standard error for all responses. Life stage is indicated on the x-axis as hours post fertilization (hpf; eggs=brown, embryos=yellow, larvae=cyan, metamorphosed polyps=green, attached recruits=pink). P-value of Kruskal Wallis tests (KW) or analysis of variance tests (ANOVA) displayed in each panel. Letters indicate significance of post hoc comparisons with shared letters indicating no significant difference.

Size-normalized respiration rates also changed across ontogeny (Kruskal-Wallis, χ^2^(5,91)=55.68, P<0.001), decreasing steadily from 65 hpf embryos to the lowest observed value at 231 hpf in larvae (Fig 2B). Respiratory demand then increased sharply in metamorphosed polyps, at 183-231 hpf, with rates significantly higher than swimming larvae of the same age (Fig 2B). The largest difference was seen at 183 hpf with respiration rates 83% higher than swimming larvae of the same age (Fig 2B). Similarly, size-normalized gross photosynthesis rates exhibited a significant decrease across development (Kruskal-Wallis, χ^2^(5,91)=34.98, P<0.001), with the lowest photosynthesis rates at 231 hpf in both swimming larvae and metamorphosed polyps (Fig 2C). Comparison of the ratio of oxygen production to oxygen demand was calculated as P:R ratio, where values >1 indicates a surplus of oxygen produced. We found that P:R ratios changed significantly across development (Kruskal-Wallis, χ^2^(5,91)=38.91, P<0.001), characterized by an initial increase from embryos at 65 hpf to larvae at 183 hpf, peaking at 2.71 ± 0.18 (mean ± standard error of mean; Fig 2D). After this point in development, P:R ratios decreased significantly (Fig 2D), reaching a minimum (1.19 ± 0.16) in metamorphosed polyps (183-231 hpf), driven by elevated rates of respiratory demand at this stage (Fig 2B).

Endosymbiont populations increased across development when calculated as total cells per individual (ANOVA, SS=1.02e8, F(9,30)=25.06, P<0.001), peaking in the oldest life stages characterized (larvae and metamorphosed polyps at 231 hpf) (Fig S1). Between the earliest larval stage and the latest metamorphosed polyp stage, there was a 138% increase in cell densities per individual (post hoc P<0.001). Because the size of life stages changed across development and cells per individual correlated positively with size (Spearman’s correlation, S=4876, rho=0.543, P<0.001), we further calculated cells per unit volume (cells mm^3^). There was a significant change in size-normalized cell density between time points (ANOVA, SS=5.29e10, F(9,30)=21.28, P<0.001) (Fig 2E). Size-normalized cell densities were highest between 65-93 hpf (post hoc P<0.05). Densities at 65 hpf were 300% higher than density at the previous stage of 38 hpf and population density decreased by 43% from 93 hpf to 163 hpf (Fig 2E). There was a trend for increased cell densities within attached recruit stages (183, 231, and 255 hpf), but this was not statistically significant (Kruskal-Wallis, χ^2^(2)=5.56, *P*=0.062; Fig 2F). Due to different units of size normalization, we cannot directly compare symbiont densities between attached recruits and earlier stages.

### B. Symbiotic and microbial community composition across development

Our samples contained a mixture of *Cladocopium* sp. (ITS2 Profile C31.C17d.C21.C21ac.C31a; ^32^) (56.0 ± 0.42%) and *Durusdinium glynnii* (ITS2 Profile D1.D4.D6.D1ab.D17d; ^33^) (44.0 ± 0.42%). Symbiont community similarity at the level of Defining Intragenomic Variants (DIVs ^34^) significantly changed across development (PERMANOVA, SS=0.09, DF=9, R^2^=0.50, F=2.97, P=0.001; Fig S2AB) but there was no change in alpha diversity (Shannon and Inverse Shannon metrics, P>0.05 for all metrics). We calculated the ratio of relative abundance of *Cladocopium* sp. and *Durusdinium glynnii* (C:D ratio) across development. We found a significant effect of time point on C:D ratios (one-way ANOVA, SS=0.38, DF=9, F=3.06, P=0.011) (Fig S2C); however, there was only a significant difference in C:D ratios found between eggs (1 hpf) and one larval (231 hpf stage (increased C:D ratio; post hoc P=0.025; Fig S2C) with no other significant pairwise differences (P>0.05; Fig S2C). Relative abundance of C31 was elevated at 231 hpf in larvae compared to eggs at 1 hpf, specifically (Fig S2B).

We further investigated bacterial community composition in *M. capitata* early life stages. However, due to lower than anticipated sequencing depth in our samples, we constrained our analysis at the taxonomic level of genus, when available, or family and above. Bacterial communities varied significantly across development (PERMANOVA, SS=4.879, R^2^=0.643, F(7,17)=4.378, P=0.001) (Fig S3A), and alpha diversity peaked during metamorphosis and recruitment (one-way ANOVA; Shannon: SS=20.247, F(7,17)=22.31, P<0.001; Inverse Simpson: SS=1.69e4, F(7,17)=17.46, P<0.001) (Fig S3B). Specifically, we found that Alteromonadaceae (*Alteromonas* sp.), *Vibrio* sp., and Pseudoalteromonadaceae (*Psychrosphaera* sp.) peaked early in development followed by Oceanospiriliaceae (unclassified) increasing in larval stages (Fig S3CD). In larvae and recruits, the relative abundance of Rhodobacteraceae and Flavobacteriaceae also increased (Fig S3CD).

### C. Ontogenetic shifts in gene expression

There were strong shifts in gene expression across development (PERMANOVA; SS=3.19e5, R^2^=0.75, F(9,28)=9.36, P=0.001), as characterized from all genes that were expressed and passed filtering thresholds (*n* = 11,475 genes). Gene expression displayed three clear ontogenetic groupings in early (eggs at 1 hpf through embryos at 38 hpf), mid (embryos at 65 hpf through larvae at 163 hpf) and late developmental stages (larvae at 183 hpf through attached recruits at 255 hpf) (Fig 3A). 66% of the variance in gene expression was explained by the separation of early developmental stages (PC1; Fig 3A) with 16% of the variance explained by the separation between mid and late developmental groupings (PC2; Fig 3A). Of the 11,475 genes expressed, 9,181 were detected as differentially expressed (DEG) through likelihood ratio tests (Fig S4). Expression patterns of the DEGs across development (i.e., clusters) were then detected through hierarchical clustering analyses with 64 total clusters identified that contained at least 15 genes (Fig S5; Table S2). 9,072 out of the 9,181 DEGs (98.8% of DEGs) were assigned to a cluster. These clusters all fell within one of three main patterns with peak expression in either early (1-38 hpf), mid (65-163 hpf), or late (183-255 hpf) developmental groupings (Table S2) and were combined for analyses of each of the three developmental patterns (Fig 3B). In total, 2,679 genes were identified as peaking in expression in early development, 3,192 peaking in mid-development, and 3,201 peaking in late development (Fig 3B).

**Fig 3.**
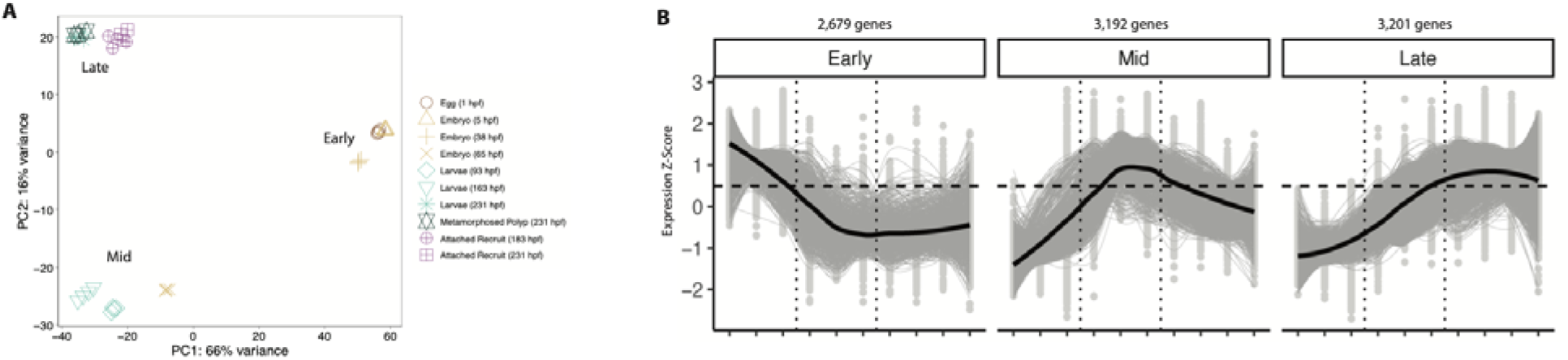
Shifts in gene expression across coral early life history stages. (A) Principal components analysis (PCA) of gene expression across *M. capitata* ontogeny (hours post-fertilization and life stage indicated by shape) in eggs (brown), embryos (yellow), larvae (cyan), metamorphosed polyps (green) and attached recruits (pink). Developmental patterns in gene expression are labeled as early (1-38 hpf), mid (65-163 hpf) and late (183-255 hpf) stages. (B) Gene expression (z-score) of DEGs peaking in early (1-38 hpf), mid (65-163 hpf) or late (183-255 hpf) developmental time points (x-axis). The number of DEGs within each developmental pattern are displayed above each plot. Gray lines indicate expression across time points for individual genes with black lines indicating the mean of all DEGs in the respective group. Dotted vertical lines indicate 38 hpf and 163 hpf separating early, mid, and late developmental phases.

There were 386 GO terms enriched in early development (eggs at 1 hpf through embryos at 38 hpf) that included functions involved in embryonic development including cell growth, cell cycle, and DNA and RNA processing cell cycle, nucleic acid metabolic process, mRNA processing, chromosome organization, RNA processing and localization, signal transduction, cytoskeleton organization, developmental process, cell division, and methylation and demethylation (Fig S6). Early development gene expression was dominated by GO terms involved in development and not by functions related to energetics or metabolism (Table S3).

In mid-development (embryos at 65 hpf through larvae at 163 hpf), there were 27 enriched GO terms (Table S4). These terms included terms related to morphological development and swimming behavior including cilium movement and cilium-dependent cell motility (Fig 4A). A full list of GO terms enriched and statistical results in mid-development are provided in Table S4. There were also terms related to energy and central metabolism including ATP metabolic processes, organonitrogen compound biosynthetic process, oxidative phosphorylation, translation and protein targeting (Fig 4A).

**Fig 4.**
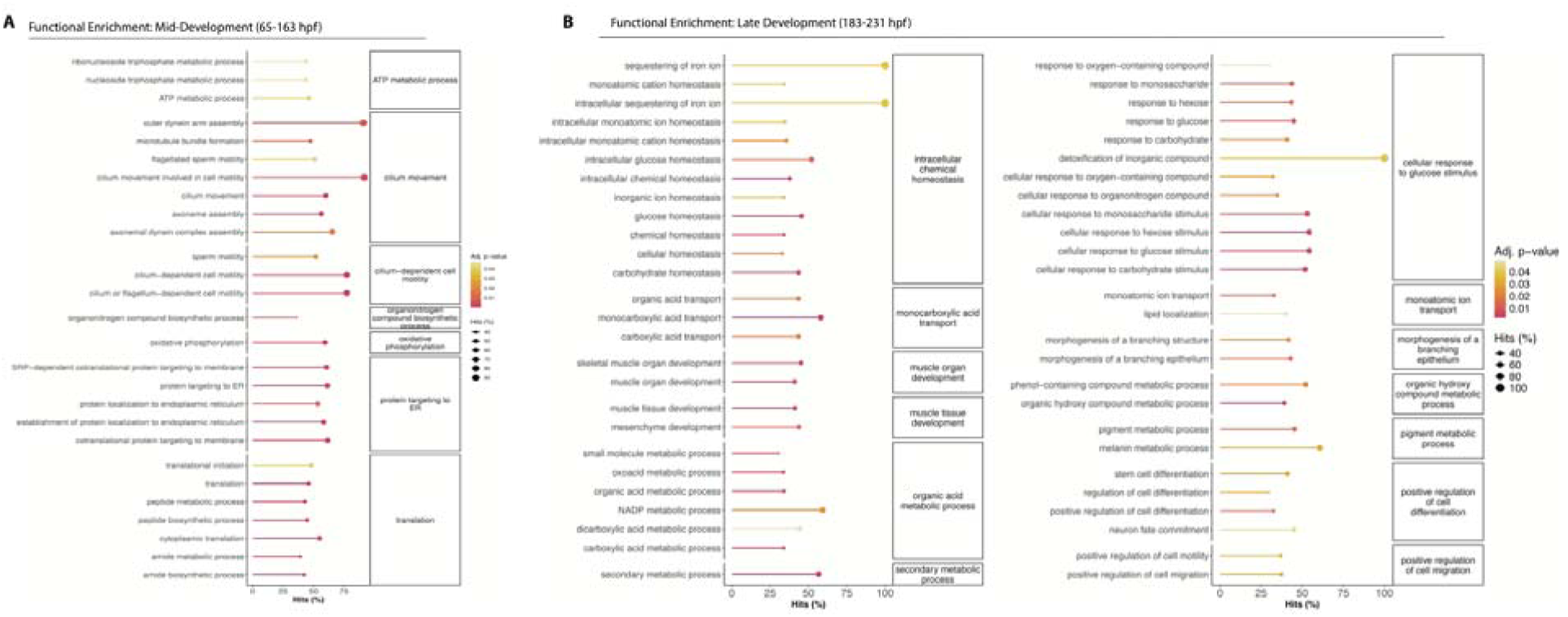
Functional enrichment of differentially expressed genes in mid and late development. Lollipop plots of the enrichment of significant (adjusted P<0.05) gene ontolog (GO) terms (Biological Process; BP) in (A) mid and (B) late development stages. x-axis and dot size indicate % hits (percentage of the number of DEGs in the respective GO category compared to total number of genes included in that category) with color indicating P-value. Parent term is indicated in panels on the right and individual GO terms on the left.

Gene expression in late development (183-255 hpf including larvae, metamorphosed polyps, and recruits) was also enriched for terms involved in morphological development including muscle tissue and organ development, morphogenesis of a branching structure, and cell differentiation including stem cell differentiation and neuron fate commitment (Fig 4B). A full list of GO terms enriched and statistical results in late development are provided in Table S5. There was also enrichment of GO terms related to intracellular chemical and ion homeostasis, including sequestration of iron ions, intracellular cation homeostasis, and inorganic ion homeostasis (Fig 4B). In addition, there was enrichment of terms related to ion transport, detoxification of inorganic compounds, and response to oxygen-containing compounds (Fig 4B). There were several enriched terms directly related to glucose and carbohydrate availability and metabolism. These terms included glucose and carbohydrate intracellular homeostasis, response to glucose, hexose, and carbohydrates, and cellular response to glucose, hexose, and carbohydrate stimuli (Fig 4B). Central metabolism terms including organic acid, oxoacid, carboxylic acid, and NADP metabolic processes were enriched, as was organic and carboxylic acid transport and lipid localization (Fig 4B).

### D. Holobiont metabolomic responses across ontogeny

There were significant shifts in holobiont metabolomic responses across ontogeny characterized as changes in metabolite pool size from 183 identified compounds (PERMANOVA; SS=5361.80, R^2^=0.64, F(12,46)=5.13, P=0.001). Unsupervised PCA visualizations showed shifts in metabolomic responses across time points (Fig 5A) with 34.7% of variance on PC1 describing shifts across developmental time and 12.09% of variance on PC2 primarily describing separation of attached recruits from other life stages (Fig 5A). Due to the significant shifts in metabolites seen in unsupervised analyses, we proceeded with supervised PLS-DA analysis (Fig 5B) and variables of importance on projection (VIP) identification to determine metabolites that significantly contribute to variation across time points. PLS-DA analyses showed that 40% of variance was explained on X-variate 1, describing shifts across developmental time, with X-variate 2 explaining 12% of variance driven primarily by separation of eggs and attached recruits from other life stages (Fig 5B).

**Fig 5.**
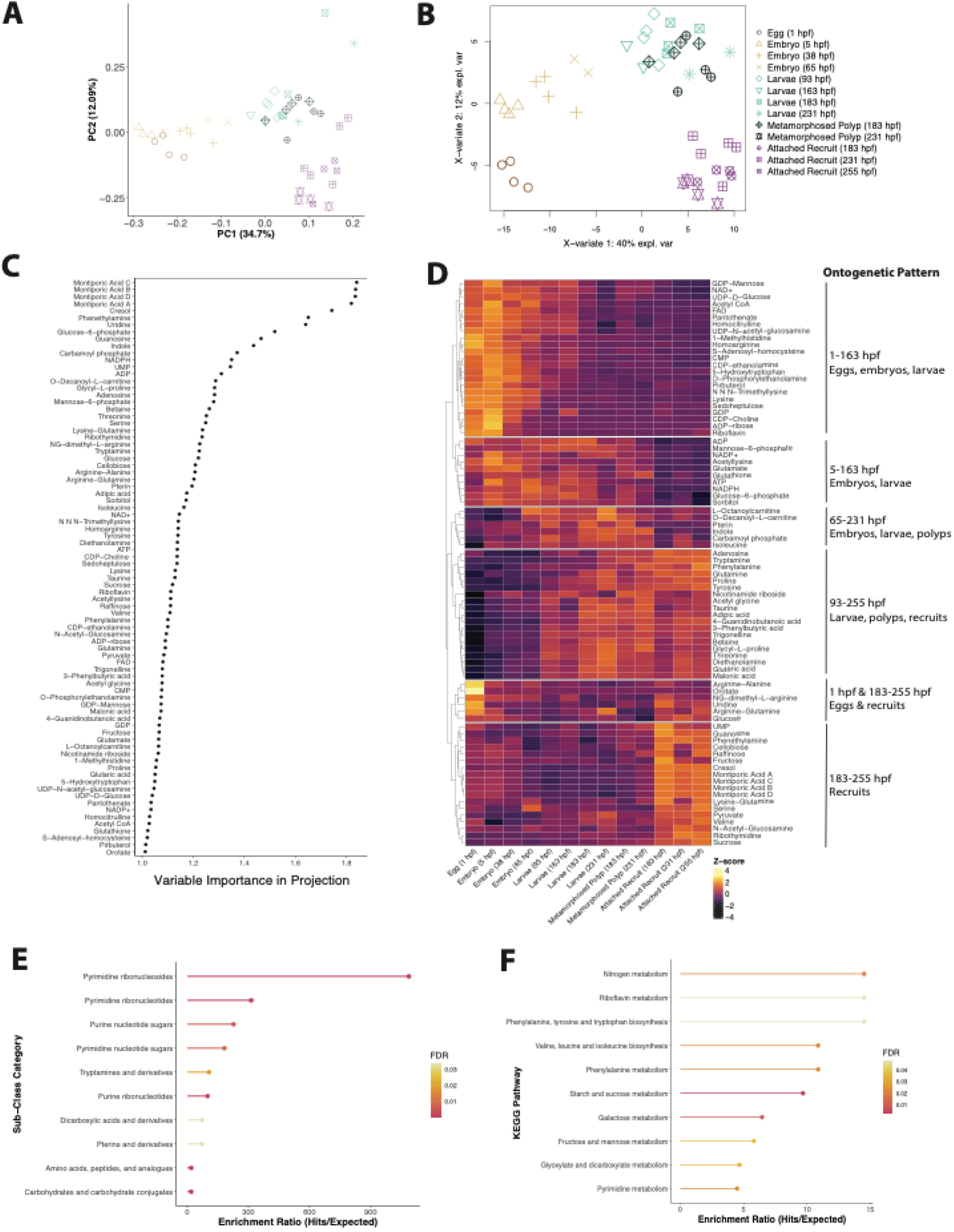
Metabolomic transitions across *Montipora capitata* ontogeny. (A) Unsupervised principal components analysis (PCA) of multivariate metabolite concentrations (normalized pool size) across *M. capitata* ontogeny (time point indicated by shape) in eggs (brown), embryos (yellow), larvae (cyan), metamorphosed polyps (green) and attached recruits (pink). (B) Supervised partial least squares discriminant analysis (PLSDA) of multivariate metabolite concentrations. (C) Variable importance in projection values for metabolites that are statistically significant in discriminating between time points (VIP _≥_1). (D) Heatmap of expression (z-score) of VIP metabolites (82 total VIPs) across time points (x-axis). Orange indicates increased concentration of a metabolite relative to the mean of all samples, with purple indicating reduced concentration. Annotations indicate ontogenetic patterns as developmental periods in which metabolite concentration peaks. (E) Sub-class metabolite enrichment analysis conducted in MetaboAnalyst showing significantly (P<0.05) enriched classes in VIP metabolites. X-axis indicates enrichment ratio (number of VIP metabolites in the respective pathway / number of expected metabolites in the respective pathway) with color indicating significance of adjusted P-values. (F) KEGG pathway enrichment analysis conducted in MetaboAnalyst showing significantly (P<0.05) enriched pathways in VIP metabolites.

We identified 82 VIP metabolites with VIP values ≥ 1 that significantly contribute to separation between time points (45% of all metabolites; Table S6; Fig S7; Fig 5C). Four Montiporic Acids displaying the highest VIP values (ranging from 1.82 to 1.84), all peaking in attached recruits (Table S6; Fig 5D). Pool sizes of VIP metabolites changed across ontogeny, with groups of metabolites peaking during distinct developmental periods (Fig 5D; Table S6). Specifically, there were 23 metabolites peaking in early development in egg, embryo, and larval stages from 1-163 hpf. These metabolites include purine and pyrimidine nucleotide sugars and amino acids (e.g., FAD). There were 10 metabolites also peaking in early development in embryos and larvae from 5-163 hpf, including ATP and ADP, carbohydrates (e.g., glucose-6-phosphate), and amino acids (e.g., glutathione, glutamate). In mid-development, there were 6 metabolites with elevated pool size, including organic phosphoric acids (e.g., carbamoyl phosphate), fatty acid esters (e.g., carnitines), and indole. As development progressed into larval, polyp, and recruit stages between 93-255 hpf, metabolites including amino acids (e.g., betaine, glutamine), dicarboxylic acids, phenylpropanoic acids, and organosulfonic acids increased in concentration. In attached recruit stages (183-255 hpf), there was a peak in several carbohydrates (e.g., cellobiose, sucrose, fructose), pyruvate), and amino acids. Interestingly, a set of metabolites peaked in both eggs (1 hpf) and attached recruits (183-255 hpf) including glucose, dipeptides (arginine-alanine, arginine-glutamine), and ribonucleosides.

To examine metabolic pathways that shift across development, we conducted enrichment analyses of metabolite classes (Table S7) and KEGG pathways (Table S8) of all VIP metabolites. At the metabolite class level, there was significant enrichment of pyrimidine ribonucleosides (p<0.001) and nucleotides (p=0.007) as well as purine ribonucleotides (p<0.001), and purine (p=0.005) and pyrimidine nucleotide sugars (p=0.007) in metabolites that discriminate between developmental stages (Fig 5E). Tryptamines (p=0.019), dicarboxylic acids (p=0.034), pterins (p=0.034) and their derivatives were also enriched in VIP metabolites along with carbohydrates and conjugates (p<0.001) and amino acids, peptides, and analogous (p<0.001) (Fig 5E). KEGG pathways involved in carbohydrate metabolism including starch and sucrose (p=0.001), galactose (p=0.009), fructose and mannose (p=0.036), and riboflavin metabolism (p=0.049) were enriched in VIP metabolites along with glyoxylate and dicarboxylate metabolism (Fig 5F). Nitrogen (p=0.019) and amino acid metabolism (e.g., phenylalanine (p=0.022), valine, leucine, and isoleucine (p=0.022), and phenylalanine, tyrosine, and tryptophan metabolism (p=0.049)) were also enriched in metabolites that changed across development (Fig 5F).

Finally, we examined changes in pool size of four VIP metabolites of interest related to symbiotic nutritional exchange including glucose (a primary photosynthate) ^21,35–37^, glutamine and glutamate (metabolites involved in ammonium assimilation) ^9,31,38,39^, and arginine-glutamine (a dipeptide involved in nitrogen sequestration) ^31,40^. Glucose concentrations varied significantly across development (ANOVA, SS=0.46, F(12,34)=4.16, P<0.001), peaking in attached recruits at 231 hpf with lowest levels see in metamorphosed polyps at 231 hpf (post hoc P<0.05; Fig 6A). Glutamine exhibited significant increases across development (ANOVA, SS=3.40, F(12,34)=35.01, P<0.001). Glutamine concentrations were lowest in eggs and reached a peak in attached recruits at 183-255 hpf (post hoc P<0.05; Fig 6B). Glutamate exhibited the opposite pattern across development, decreasing significantly (ANOVA, SS=0.54, F(12,34)=7.96, P<0.001) from eggs and embryos to attached recruits (Fig 6C). Arginine-glutamine concentrations also varied significantly (ANOVA, SS=2.17, F(12,34)=17.50, P<0.001) with a peak in eggs, larvae (231 hpf), and attached recruits (183-255 hpf) (Fig 6D).

**Fig 6.**
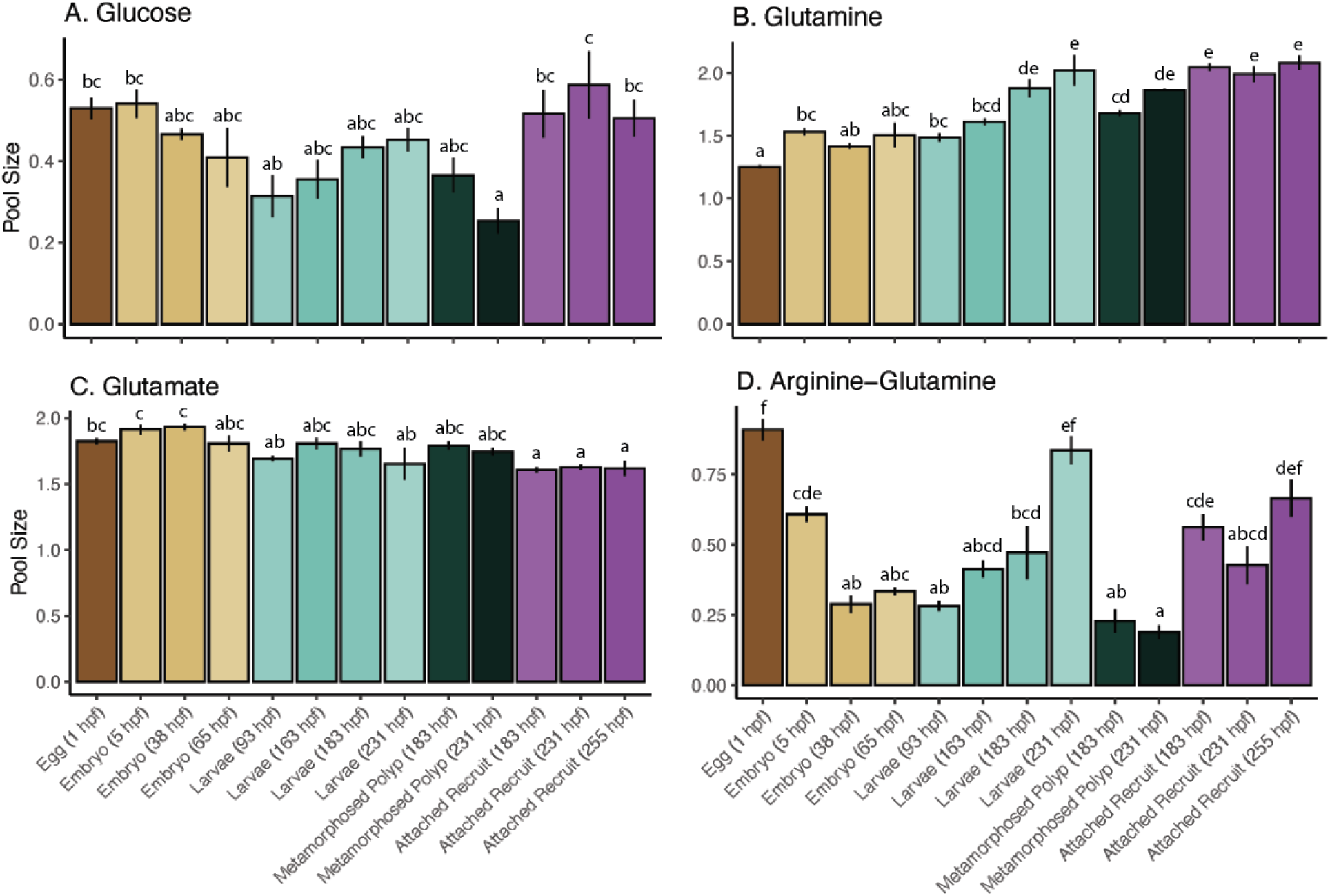
Changes in key metabolite concentrations related to nutritional exchange and nitrogen cycling across *Montipora capitata* development. Pool size (log-transformed median-normalized) of four metabolites of interest: (A) glucose, (B) glutamine, (C) glutamate, and (D) arginine-glutamine. Data are represented as mean ± standard error for all metabolites. Life stage is indicated on the x-axis as hours post fertilization (hpf) (eggs=brown, embryos=yellow, larvae=cyan, metamorphosed polyps=green, attached recruits=pink). Letters indicate significance of post hoc comparisons with shared letters indicating no significant difference.

## Discussion

The results of this study provide an extensive organismal and multi-omic view of metabolism and symbiotic interactions across multiple early life history stages of a vertically transmitting coral, advancing understanding of metabolic shifts during development in a foundational nutritional symbiosis. Critically, the results of our study detail mechanisms by which corals undergo extensive metabolic changes during development and that both symbiont-derived and stored energy reserves are important to support energetic demands (Fig 7). We found that although nutritional exchange may begin in larval stages in this species ^31^, a strong metabolic response to carbohydrate photosynthates was not seen until after metamorphosis (Fig 7). In addition, we found that lipid metabolism was important in early development with clear evidence of increased importance of symbiont-derived nutrition after metamorphosis (Fig 7). Collectively, our integrative approach clarifies and deepens understanding of ontogenetic metabolic shifts and highlights the large molecular and metabolic reorganization that occurs during coral early life history. Because the metabolism of symbiont-derived nutrition increases during sensitive post-settlement stages, symbiotic disruption under climate change driven environmental stress has the potential to reduce energetic capacity and, consequently, successful coral reef recruitment and persistence.

**Fig 7.**
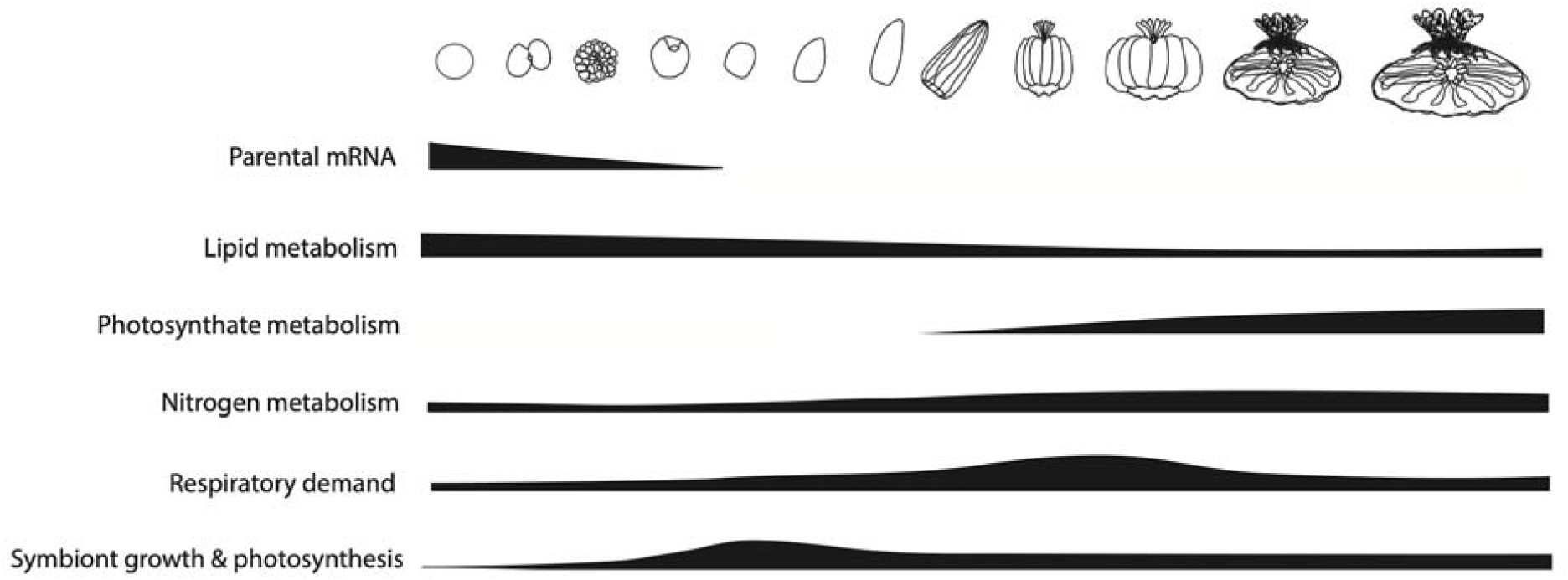
Developmental shifts in metabolism across *Montipora capitata* ontogeny observed in this study. Thicker bars indicate qualitative representation of increased relative contribution of each metabolic pathway or energy source to development.

### A. Early developmental energy demands are supported by parentally provisioned reserves

Stored parental reserves are an important source of nutrition in early stages of coral development. Coral offspring are lecithotrophic (i.e., feed on maternally provisioned egg yolk) and under normal environmental conditions are provisioned with stored energy reserves sufficient to complete development until symbiotic nutrition and/or heterotrophic feeding provide necessary energy for growth and survival ^23,26,28,41,42^. In early development, embryos undergo a reorganization of tissues and cellular structures, gastrulation and differentiation of germ layers, and rapid cell proliferation ^43–45^. These processes require maternal and paternal transcripts, structural molecules, and metabolic building blocks (e.g., glycogen, amino acids, lipids) to support cell functions that are provisioned to differing extents to the eggs from the parents ^46–48^. The total amount and rate at which these lipids are metabolized can determine larval pelagic duration and energy available to meet the demands of development, metamorphosis, settlement, and survival ^49–51^.

We found evidence of metabolism of parentally provisioned lipids in early embryos (1-38 hpf). Specifically, there were increased concentrations of intermediates and cofactors involved in lipid and stored energy metabolism in early development (e.g., acetyl CoA, lysine, pantothenate). Glyoxylate metabolism was one of the enriched metabolic pathways that changed across development. The glyoxylate pathway generates acetyl CoA from lipid stores when carbohydrate availability is limited and may facilitate stored reserve metabolism during stress and bleaching ^52–55^. Furthermore, there were increased concentrations of cofactors involved in fatty acid and central metabolism including pantothenate, riboflavin, and lysine and precursors of glycerol storage (e.g., UDP-Glucose). Our results support that metabolism of stored lipids contribute to the energetic needs of early embryonic development.

Lipid metabolism also played a role in supporting development throughout larval stages. During larval stages, there were increases in carnitine metabolites (i.e., O−Decanoyl−L−carnitine and L−Octanoylcarnitine), which transport fatty acids during lipid metabolism, suggesting that larvae were continuing to metabolize lipid stores, as demonstrated in previous research ^24–26^. However, there was no enrichment of lipid metabolism processes found in our gene expression analysis and in combination with our observation of decreased concentrations of lipid metabolism cofactors (e.g., acetyl CoA, lysine, pantothenate, riboflavin) supports the finding that stored lipid metabolism decreases in contribution as development progresses. Parental provisioning is critical for early embryonic development in our study and underscores the importance of characterizing how environmental stressors may compromise gamete provisioning, especially during reproductive events following mass bleaching events ^48,56^.

### B. Shifts in nitrogen metabolism across ontogeny support nutritional requirements and symbiotic relationships

Nitrogen cycling is a central function in the coral-algal symbiosis and is a driver of symbiotic stability and productivity ^39,57–59^. We found that nitrogen metabolism changed significantly across development to support nutritional demands and maintain a stable symbiotic relationship. Nitrogen availability can be modulated by both the host and symbiont through assimilation of free ammonium by each partner through the glutamine synthetase-glutamine oxoglutarate aminotransferase/glutamate synthase pathways (GS-GOGAT) ^8,38,60–62^. Through these pathways, ammonium is assimilated into amino acids through sequential action of glutamine synthetase (GS; produces glutamine from glutamate and ammonium) and glutamine oxoglutarate aminotransferase/glutamate synthase (GOGAT; produces two molecules of glutamate from glutamine and α-ketoglutarate), which generates amino acids that can be used for a wide variety of cellular functions and metabolism ^38,40,63^.

Signatures of increased ammonium metabolism in our study first appear in mid-developmental stages with significant shifts in the concentration of glutamine, glutamate, and arginine-glutamine dipeptides in larvae. Our metabolomic results revealed a consistent increase in glutamine pools with concurrent decrease in glutamate pools across development, indicating activity of the glutamine synthase pathway to assimilate ammonium in the holobiont through production of glutamine ^38,64^. This was likely driven by nitrogen uptake in the algal symbiont community, because ammonium assimilation by the symbiont can account for 14-23 times greater ammonium assimilation than host assimilation in adult corals ^38^ and there was no enrichment of nitrogen metabolism functions in host gene expression. Importantly, we observed increased symbiont densities between 65-93 hpf in larvae as well as increased photosynthesis between 93-183 hpf, corresponding with this period of elevated ammonium assimilation by the algal symbiont in larval stages and further suggesting symbiont proliferation is a nitrogen-limited response ^65^.

Other mechanisms of nitrogen mechanisms, such as production arginine-glutamine dipeptides have been hypothesized to play a role in nitrogen sequestration by the coral host in adult corals ^40^ and in *Montipora capitata* larvae ^31^. Interestingly, there were three distinct peaks in arginine-glutamine dipeptide concentrations across development - first in eggs (1 hpf), then in larvae (231 hpf), and finally in attached recruits (183-255 hf). The high production of arginine-glutamine in our study during larval and recruit stages suggests that the coral host is actively sequestering nitrogen to reduce nitrogen availability to the symbiont, as demonstrated previously in *M. capitata* larvae ^31^. Nitrogen limitation of the symbiont population reduces the rate of population growth and can favor the translocation of excess photosynthates from the symbiont to the host ^2,8,39,57,58,60,66^. Recent research in the coral *Stylophora pistillata*, the sea anemone *Exaiptasia diaphana*, and jellyfish *Cassiopea andromeda* have demonstrated that passive feedback loops between carbon and nitrogen availability are crucial for a stable symbiosis and disruption in these feedback loops can lead to dysbiosis ^39,58,59^.

We suggest that alongside ammonium assimilation via the GS pathway, the coral host performs nitrogen sequestration via the production of arginine-glutamine dipeptides during early development to stabilize symbiont growth and translocation. We found that symbiont population densities decreased by 43% after 93 hpf and were stable from 163 hpf onward. Similarly, photosynthesis rates increased a few hours later at 183 hpf and remained stable further into development. This stabilization in cell densities and photosynthetic rates supports the interpretation of regulation of nitrogen availability to symbiont populations by the host ^8,66,67^. Alongside metabolomic evidence of nitrogen metabolism, we documented increased gene expression enrichment for host response to organonitrogen compounds and monatomic ion transport, which could indicate increased transport of ions such as ammonium and/or bicarbonate ^68–70^. Also increased in mid-development, were carbamoyl phosphate concentrations which are involved in the urea cycle to remove nitrogen through the production of urea ^31,40,71^. In a previous study in *M. capitata* larvae, the host used ammonium assimilation and nitrogen sequestration via the urea cycle and dipeptide production as a mechanism to limit nitrogen availability to the symbiont and maintain stable symbiont populations under thermal stress ^31^. We suggest that these mechanisms continue throughout metamorphosis and settlement to maintain stable symbiotic relationships and control algal symbiont population growth.

In early development, however, arginine-glutamine dipeptides may play a different role by being used to meet nutritional requirements. Arginine-glutamine was elevated in eggs along with enriched gene expression of nitrogen metabolism, suggesting that parentally provisioned dipeptides may play different roles depending on developmental stage and may provide a source of nitrogen for developing embryos. Arginine-glutamine is more soluble than either amino acid individually ^72^ and may therefore be a stable mechanism for providing a source of nitrogen to gametes in corals. Specifically, the catabolism of arginine-glutamine may provide a source of glutamine for catabolic glutamate dehydrogenase activity that can generate a-ketoglutarate for metabolism through the tricarboxylic acid cycle ^39^. Recent work has demonstrated that heterotrophically-sourced nitrogen is indeed packaged into coral gametes ^73^ and we suggest that dipeptides may be one form of provisioned nitrogen. Future work should examine the role of dipeptides in supporting the nutritional needs of developing embryos and how the role of this metabolite changes across development.

Throughout this developmental time series, our samples contained a mixture of *Cladocopium* sp. and *Durusdinium* sp., which is characteristic of this species in KāneC:ohe Bay, HawaiC:i ^74,75^. Stability in Symbiodiniaceae communities suggests that the nutritional and metabolic changes documented in this study are not driven by changes in symbiont identity and thus function. However, as environmental conditions continue to change, the thermal tolerance advantages of hosting *D. glynnii* previously documented in adult *M. capitata* ^76,77^ could manifest in a shift to a greater abundance of *D. glynnii* than *Cladocopium sp.* provisioned to offspring ^78^. Interestingly, *Acropora tumida* recruits (approx. 1.5 months old) exhibit higher infection rates of *Durusdinium* symbionts under elevated temperature, but with no clear fitness advantages 31 days after infection ^79^. Since *D. glynnii* translocate lower amounts of photosynthates than *Cladocopium sp.* in *M. capitata* ^80^, this could result in detrimental impacts on nutritional availability for offspring in the future and tradeoffs that can result in reduced growth and reproduction at later stages and should be examined in additional work.

Bacterial communities play important roles in nutritional cycling ^57,81,82^ and acquisition of micronutrients ^83^, and can act as nutritional mediators between host and symbiont ^84,85^. For example, larvae of the brooding coral *Pocillopora acuta* transfer bacterially derived nitrogen to their symbiont populations ^84^. Shifts in bacterial taxa indicate potential changes in microbial nutritional relationships in early life history. We found that Alteromonadaceae (*Alteromonas* sp.), Pseudoalteromonadaceae (*Psychrosphaera* sp.) were highest in relative abundance in early development, with Oceanospiriliaceae peaking in larvae and Rhodobacteraceae and Flavobacteriaceae highest in late development. These taxa play important roles in carbon ^86^, nitrogen ^84^, sulfur (e.g., DMSP/DMS ^87^), and micronutrient cycling ^83^ along with the production of antimicrobial compounds ^88^. Further, taxa in Rhodobacteraceae are associated with symbiont communities and may signal a shift in the microbial communities associated with symbionts across development ^89^. Therefore, we hypothesize that our observations of an increase in bacterial diversity could confer increased nutritional functional diversity and capacity for nutrient cycling as corals age ^90,91^. Although our limited sequencing depth of microbial communities prevented us from making detailed functional conclusions on shifts in bacterial communities, the increases in microbial diversity across ontogeny are compelling and are deserving of future research to understand the functional and nutritional role of bacterial associations during coral development. To test this hypothesis, quantitative measures of nutrient cycling between bacterial communities and the coral host should be completed in future studies with approaches such as metatranscriptomics, DNA-stable isotope probing (e.g., DNA-SIP), as well as metabolomic stable isotope probing and microscopy techniques (e.g., NanoSIMS) applied to early life stages ^92^.

### C. Photosynthate metabolism becomes increasingly important after metamorphosis

Progressing through metamorphosis and settlement is an energetically demanding process for marine taxa ^93–95^ due to rearrangement of the body plan ^96^ and initiation of attachment and biomineralization ^69,97^. The high energetic demand of metamorphosed polyps as compared to swimming larvae at the same age, indicates *M. capitata* larvae underwent high energetic stress during metamorphosis. This is also evidenced by a sharp depletion in glucose pools in metamorphosed polyps and enrichment of genes involved in oxidative phosphorylation and ATP metabolism during this period. Thus, after metamorphosis, it is critical for newly settled corals to obtain sufficient nutrition to recover from energetic stress and meet demands of calcification and growth.

Metabolism of glucose, a major photosynthate ^21,35,36^, increased after metamorphosis. Importantly, glucose was elevated in attached recruit stages following a depletion of glucose during metamorphosis, suggesting that glucose was heavily used as a source of nutrition during metamorphosis, with pools recovering to previous concentrations after settlement. Metabolomic analyses also revealed depletion of a glucose intermediate, glucose-6-phosphate, and increased production of pyruvate, a glycolysis product in late developmental stages. Additionally, there were decreases in NAD+ and NADP+, important cofactors for glycolysis and central metabolism ^98^. Together, these shifts in glycolysis metabolites and cofactors indicate increased glycolytic metabolism after metamorphosis, as more pyruvate is produced from high turnover of intermediate metabolites. Strong enrichment of carbohydrate, glucose, and hexose metabolism, stimulus, and homeostasis in gene expression of the host provide evidence that central carbon metabolism is occurring in the host, that glucose was indeed translocated from the algal symbiont, and photosynthate metabolism plays an important role in supporting nutritional requirements after settlement.

Carbon produced through photosynthesis is the primary source of carbon packaged into coral gametes ^48^. In early development, we also found that glucose supported developmental energy demands, but that this was likely not a result of active translocation from the symbiont. The earliest developmental stages in *M. capitata* are unlikely to experience active translocation from their symbionts during the first few hours of development, even though symbionts are present. This is because spawning occurs at approximately 21:45 with fertilization complete by approximately 23:00 and therefore, the first 8 hours of development are completed in the dark with no potential for photosynthetic activity. Therefore, glucose pools in the eggs were likely parentally provisioned as a source of nutrition to support early development. As climate change intensifies and coral reproduction occurs under higher temperatures, symbiotic nutritional exchange is increasingly vulnerable in both adult and larval corals. A current paradigm of the field is that energetic contribution of symbiont nutrition to metabolic demands of swimming coral larvae is negligible ^30^. Previous work examining the influence of symbiotic nutritional exchange on larval energy requirements has largely occurred in brooded larvae in vertically transmitting species (e.g., ^27,28,30^), with ample lipid reserves ^26,28,99^, thus the relative contribution of symbiotic nutrition to energy needs may be less than that of smaller, spawned larvae. Here, we propose that prior research based on vertically transmitting brooding corals has underestimated the energetic contribution of symbiont nutrition to metabolic demands in spawning species. Our observation that post-settlement stages utilize photosynthates to support energy demand, and previous observations that spawned *Acropora* larvae use lipid reserves at reduced rates if they have symbionts ^29^, suggest that symbiotic nutrition is impactful for pelagic larval dispersal potential during early development for vertically transmitting spawning species. Further, the contribution of symbiont nutrition must be considered in the context of energetic demands. For example, ^30^ suggested that the contribution of symbiotic nutrition to larval energy demands is negligible compared to adult conspecific in *Pocillopora damicornis.* However, adult corals are expending energy for expensive processes such as calcification and reproduction ^100–102^, while larvae require energy for dispersal and metamorphosis ^26,95,103^, and therefore, while symbiotic nutritional exchange may be at lower rates in larvae, this does not discount that it may be energetically meaningful for life stage specific energy demands. Comparative studies that clarify the influence of symbiont nutrition on energetic demands across life history strategies in the context of life stage specific energetic budgets will be essential for understanding trajectories of reef recruitment under climate change. Specifically, future work conducting direct tracing of translocation using stable isotope methods across multiple developmental stages would further clarify the contribution of photosynthates to energy demands during vulnerable life history transitions.

The benefits of photosynthate translocation and increase in nutritional exchange are, however, not without dangers from temperature and irradiance driven production of reactive oxygen species (ROS) as photosynthesis byproducts ^104–106^. In our study, we observed that *M. capitata* larvae had enriched gene expression including detoxification and response to oxygen-containing compounds during late development. These pathways are involved with stress and ROS response to photosynthesis byproducts and are central components of metabolism in symbiotic corals ^21,70,107,108^. Elevated expression of cellular stress response genes supports that early life stages require activation of cellular stress management to neutralize oxidative stress from symbiont metabolic activity ^104–106^. Observed response to ROS, sequestration of trace metals, and increased carbohydrate production by the symbiont during late developmental stages suggest that nutritional exchange is active during *M. capitata* development and that the host is actively investing in the management and enhancement of symbiont photosynthesis. We found enrichment of iron ion sequestration in late developmental stages, which is a critical trace metal that affects symbiont population productivity and symbiotic stability ^83,109^. In addition, there was increased production of storage carbohydrates including sucrose, fructose, raffinose, and cellobiose in late developmental stages, which can be fixed by the symbiont and translocated to the host ^110^. Previous work demonstrating that pools of starches and carbohydrates decrease when the symbiosis is compromised under bleaching conditions ^111^ further suggests that increased carbohydrate production in attached recruits in our study is a product of photosynthetic carbon fixation. Indeed, the swimming larvae of *M. capitata* have the capacity to invest in symbiotic regulation ^31^ and stress response. Yet, in a future of increasing heat wave intensity and frequency ^112^, further increasing the energetic costs of cellular stress management to maintain symbiotic relationships will further hinder settlement and developmental success _106,113._

### D. Conclusions

In this study, we documented large-scale ontogenetic shifts in metabolism and symbiotic relationships in a vertically transmitting species, *Montipora capitata*, in HawaiC:i. We found that parentally provisioned reserves support early embryonic development. During larval stages, we documented increases in symbiont population and photosynthetic capacity, with evidence that this is driven by symbiont ammonium assimilation. Following this initial symbiont proliferation, the coral host employed nutrient cycling to enhance symbiont productivity and stabilize growth via nitrogen limitation ^31,58,59^ to support high energetic costs of settlement and metamorphosis. Active symbiotic nutritional exchange and metabolism of photosynthates increased strongly after settlement, highlighting their value to the holobiont success at the most vulnerable coral life stages. The results of this study demonstrate that the symbiotic relationship is maturing during early life stages, and that symbiotic interactions and nutritional exchange play important roles in shaping metabolic responses across development. Nutritional symbioses are ubiquitous in nature, and it is critical to understand how symbiotic relationships form in foundational organisms with complex early life histories. As environmental stress continues to increase on reefs, it is critical to conduct additional research to understand how symbiotic stress may impact the energetic vulnerabilities and dispersal potential of vertically transmitting species.

## Resource availability

### Lead contact

Requests for further information and resources should be directed to and will be fulfilled by the lead contact, Ariana S. Huffmyer.

### Materials availability

This study did not generate new unique reagents.

### Data and code availability

Data and scripts are openly available on GitHub as a static release at https://github.com/AHuffmyer/EarlyLifeHistory_Energetics/releases/tag/v2.1.1 and Open Science Framework (OSF) at DOI 10.17605/<u>OSF.IO/KMU7H</u> ^114^. Raw sequence files (TagSeq, ITS2, and 16S) are stored at the NCBI Sequence Read Archive under project PRJNA900235. Raw and processed metabolomic data are archived at OSF at DOI 10.17605/<u>OSF.IO/KMU7H</u> _114._

## Acknowledgements

This research was supported by the National Science Foundation Ocean Sciences Postdoctoral Fellowship to ASH (award no. 2205966), US-Israeli Binational Science Foundation to HMP and TM (award no. BSF 2016-321), National Science Foundation Rules of Life-Epigenetics (award no. EF-1921465), and was supported by a gift of the Washington Research Foundation to the University of Washington eScience Institute. As guests, we recognize and give thanks for the land and water resources of the C:āina and the traditional owners of the land, kānaka C:ōiwi, both past and present, as well as future generations, on which this experimental work was conducted in the KāneC:ohe AhupuaC:a and the islands of HawaiC:i. We are grateful for the logistical support provided by the Coral Resilience Lab and the HawaiC:i Institute of Marine Biology. We acknowledge Margaret Schedl for RNA extraction, Craig Nelson and Shayle Matsuda for discussions on 16S bacterial analyses, Eric Chiles and Xiaoyang Su for metabolomics discussions, and members of the Putnam Lab (University of Rhode Island) and Roberts Lab (University of Washington) for feedback on earlier versions of this manuscript. We acknowledge use of the computational resources of the URI Center for Computational Research for this work.

## Author contributions

Conceptualization: ASH, HMP; Data curation: ASH; Formal analysis: ASH; Funding acquisition: ASH, HMP, TM; Investigation: ASH, KHW, DMB, ES; Methodology: ASH, KHW, HMP; Project administration: ASH, HMP; Resources: HMP; Supervision: ASH, HMP; Validation: ASH; Visualization: ASH, HMP; Writing - original draft: ASH; Writing - review & editing: ASH, KHW, DMB, ES, TM, HMP

## Author contributions

The authors declare no competing interests.

## STAR Methods

### A. Key resources table

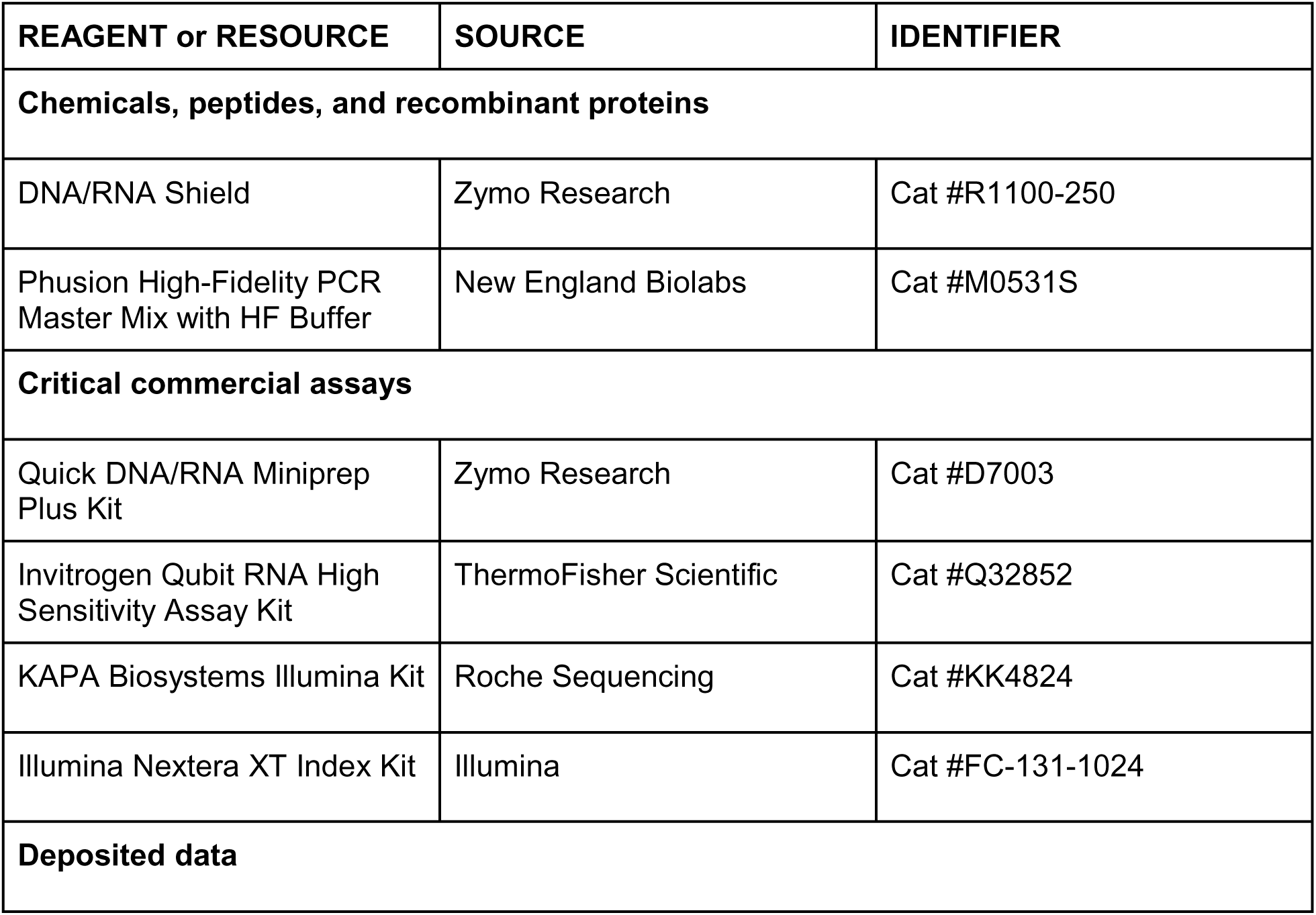
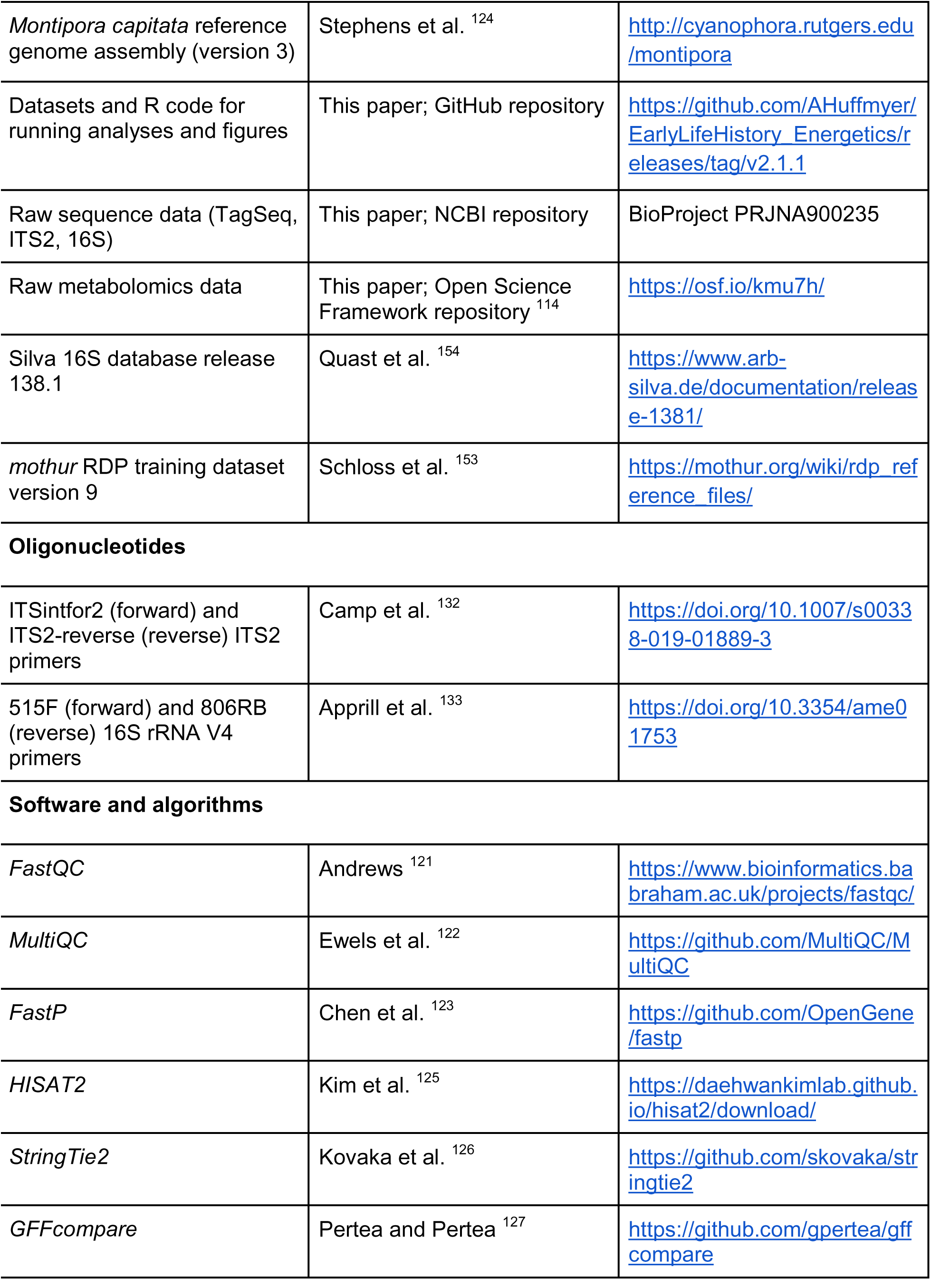
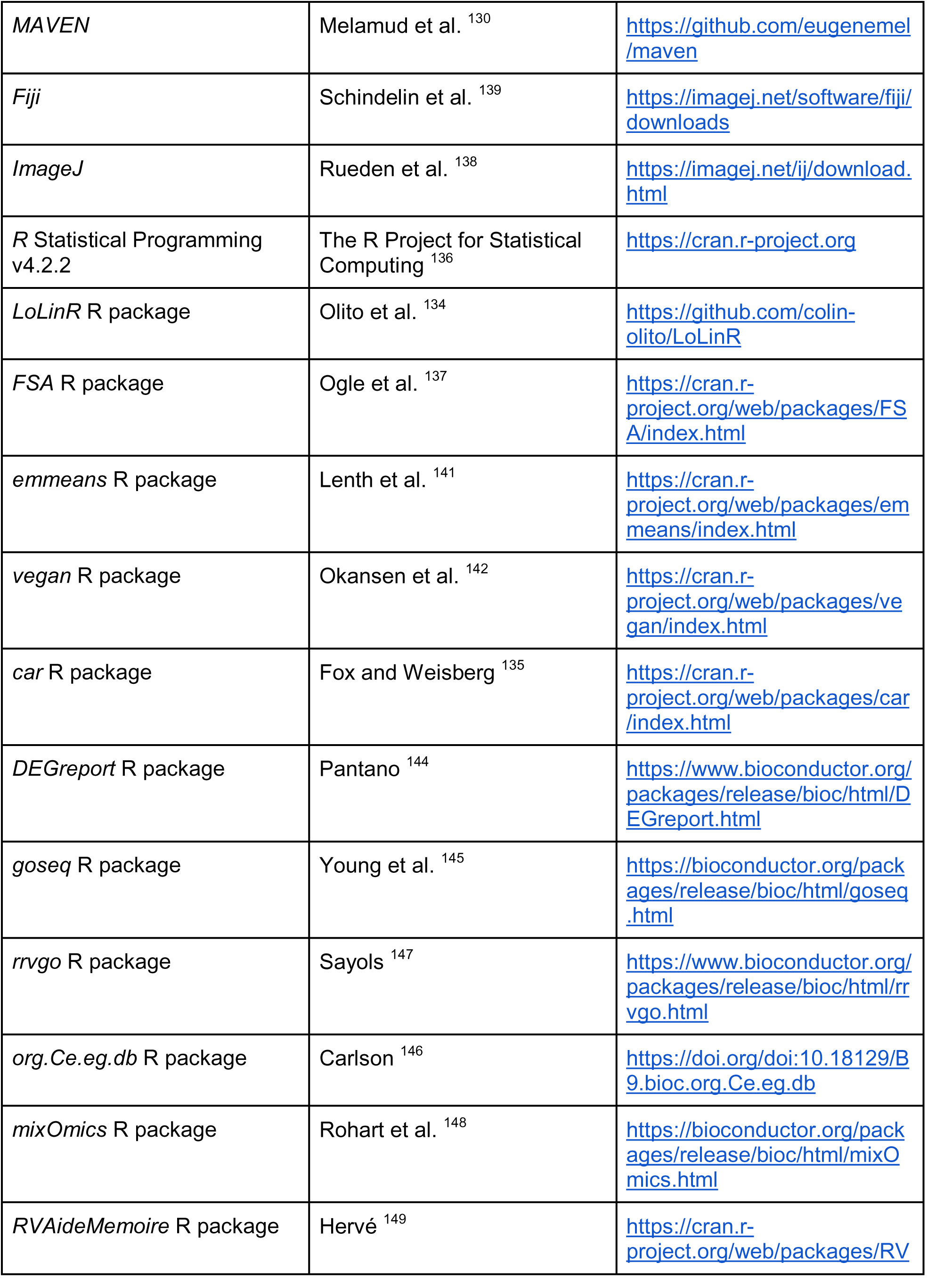
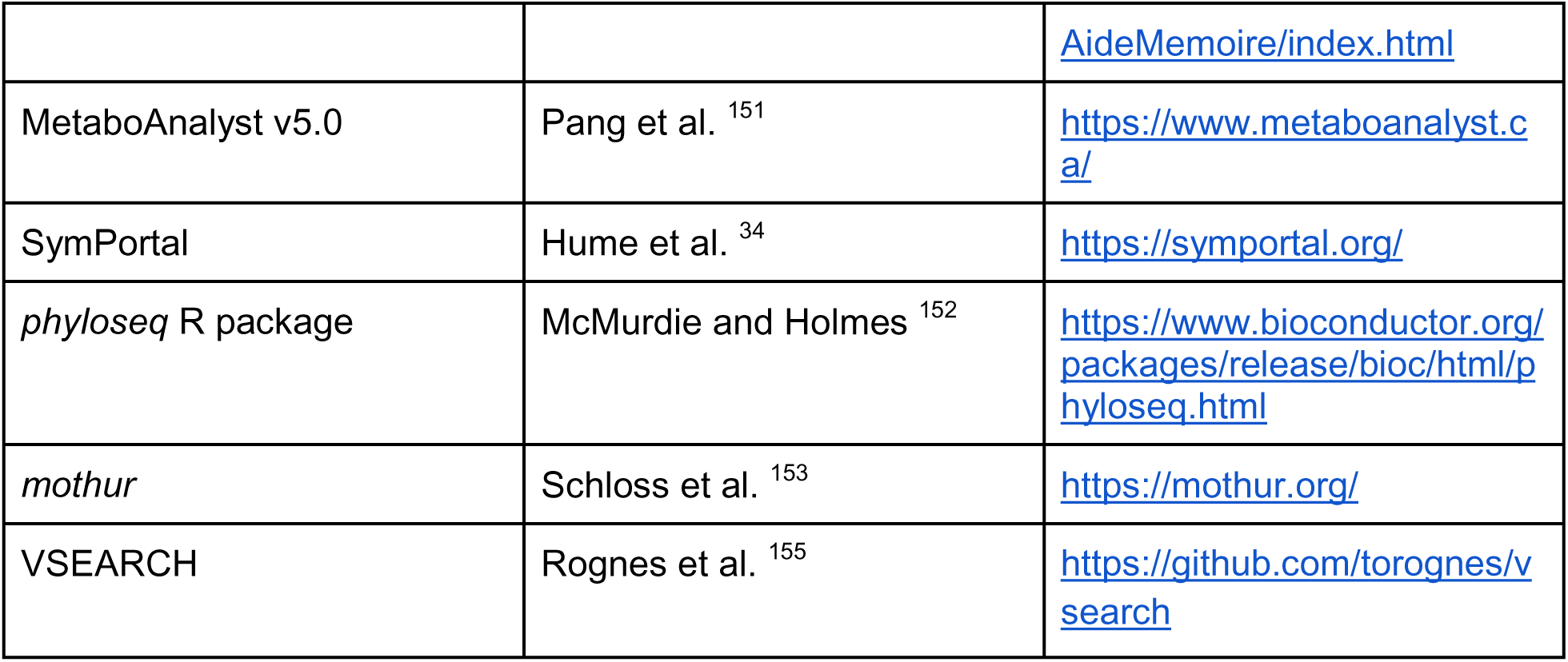

### B. Experimental model and subject details

#### i. Gamete collection and developmental rearing

*Montipora capitata* is a hermaphroditic broadcast spawning, vertically transmitting coral species ^115,116^ that is a dominant reef-builder in KāneC:ohe Bay, OC:ahu, HawaiC:i where this study was conducted (Fig 1B.1). During the July 2020 spawning period, *M. capitata* egg and sperm bundles were collected on the night (21:00) of the new moon by scooping spawned bundles from the surface of the water in KāneC:ohe Bay under the State of Hawai‘i Division of Aquatic Resources Special Activities Permit 2019-60. Egg sperm bundles were then transported to the laboratory at the HawaiC:i Institute of Marine Biology (HIMB) and approximately 3-5 mL of bundles were added to 50 mL falcon tubes filled with 0.2 µm filtered seawater (FSW) and allowed to separate and fertilize for 1 h at 27.5°C (Fig 1B.2). Following fertilization, embryos were washed with 1 µm FSW to remove sperm and then added to 20 L conical rearing vessels (*N* = 30) following methods in ^117^. Conicals were supplied with 1 µm filtered seawater (FSW) at ambient temperature at 27.5°C and 50-100 µmol photons m^-2^ s^-1^ (photosynthetic active radiation; PAR). Throughout development, embryos were gently stirred, and debris, lipid slicks, and dead material were removed every 3-6 hrs. Embryos were reared in conical vessels through development to the swimming larval stage (Fig 1B.3). At approximately 165 hours post-fertilization, larvae were transferred in 53 µm mesh-bottom plastic bins (*N* = 8; ∼ 15 cm x 30 cm) containing aragonite settlement plugs (circular surface; surface area 2.85 cm^2^) conditioned for 3 days in seawater to outdoor flow-through seawater tables for settlement following methods in ^117^. Settlement was conducted at ambient temperature (∼28°C) and light of 50-100 µmol m^-2^ s^-1^ µmol photons m^-2^ s^-1^ on a 12h:12h light:dark cycle achieved by covering bins with two layers of shade cloth. Larvae were allowed to settle in these conditions for 48 h, at which point we examined larvae for metamorphosis. Two settlement states were sampled in this study – larvae that metamorphosed and attached to the surface of aragonite plugs (Fig 1B.4) and those that remained in the water metamorphosed without attachment (i.e., metamorphosed polyps). We collected samples at 13 time points across the early life history of *M. capitata* including fertilized eggs (1 hour post-fertilization, hpf), embryos (cleavage at 5 hpf, late gastrula stage at 38 and 65 hpf), larvae (93, 163, 183, 231 hpf), metamorphosed polyps (83 and 213 hpf), and attached recruits (183, 231, and 255 hpf) (Fig 1A) for molecular, metabolomic, physiological, and Symbiodiniaceae and bacterial amplicon sequencing assays. Sample size for all response variables are presented in Table S1.

### C. Method details

#### i. Respirometry

Respiration (oxygen consumption) and photosynthesis (oxygen production) rates were measured using an Oxy-10 Sensor Dish Reader (PreSens Precision Sensing) in combination with a 24-well microplate with spot oxygen sensors (80 µL; Loligo Systems) at six stages: embryos (65 hpf), larvae (93, 183, and 231 hpf) and metamorphosed polyps (183 and 231 hpf) (Fig 1A). Six individuals were added to each 80 µL microplate well filled with 1 µm FSW (continuously aerated for 30 min prior to measurements) and sealed with glass coverslips. Sensor spots were allowed to condition for 15 min in 1 µm FSW prior to measurements. Two wells of each plate were filled with 1 µm FSW as blanks. Larvae were exposed to 10 min of dark acclimation prior to measurements. Oxygen production (“photosynthesis”) was measured as oxygen concentration (µmol L^-1^) first under light conditions for 15-20 min at ∼ 500 µmol m^-2^ s^-1^ PAR using LED aquarium lighting (AquaIllumination Hydra) with oxygen measurements collected every 15 sec. Measurements under light were conducted first to standardize light conditions prior to measurement of respiration due to light-enhanced dark respiration (LEDR) effects resulting from stimulation of respiration through photosynthesis ^118^. The lights were then turned off and LEDR oxygen consumption (“respiration”) was measured for an additional 15-20 min in the dark. Light levels during photosynthesis measurements are within the saturating irradiance for adults of this species ^119^, but PI curves were not conducted for the early life stages. Two replicate runs were conducted for embryo and larval time points (93 and 163 hpf, *n* = 10 samples per run, *n* = 6 runs). At time points where both larvae and metamorphosed polyps were analyzed (183 and 231 hpf), replicate runs were conducted in which larval and metamorphosed polyps were included in each run and randomly assigned to wells (*n* = 5 samples per time point per run, *n* = 7 runs).

#### ii. Size and symbiont density

At each time point (Fig 1A), pools of approx. 100 individuals were collected from replicate conicals (*n* = 4 conicals or bins), fixed in 4% paraformaldehyde for 24-36 h then stored in 70% ethanol at 4°C (*n* = 4 samples per time point; approx. 100 individuals per sample) for size and symbiont density measurements. Samples were digitally photographed using an OMAX Digital Microscope Camera (Model A35180U3; Kitchener, ON, CA) on a OMAX dissecting light microscope (Kitchener, ON, CA) with an OMAX 0.01 mm stage micrometer (Model A36CALM1; Kitchener, ON, CA) for calibration in each photograph. Symbiont cell density at each time point (Fig 1A) was measured by conducting replicate cell counts (*n* = 6-8 counts per sample, *n=* 4 samples per time point) of homogenized (Pro Scientific Bio-Gen PRO200 Homogenizer; Oxford, CT, USA) tissue using a hemocytometer (Hausser Scientific Improved Neubauer Hemocytometer; Horsham, PA, USA).

#### iii. Gene expression in the coral host

From all time points, except for attached recruits that were snap-frozen in liquid nitrogen, pools of 50-200 individuals were preserved in DNA/RNA Shield (Zymo Research, Irvine, CA, USA, Catalog #R1100-250). RNA and DNA extractions were performed with the Quick DNA/RNA Miniprep Plus Kit (Zymo Research, Irvine, CA, USA, Catalog #D7003) according to the manufacturer’s instructions (see detailed protocol in data repository ^114^) and stored at −80°C. RNA quantity was determined using Invitrogen Qubit™ RNA High Sensitivity Assay Kit (ThermoFisher, Waltham, MA, USA, Catalog #Q32852) and quality was assessed using 4200 TapeStation System for ribosomal bands (Agilent Technologies, Santa Clara, CA, USA). Purity was assessed using a NanoDrop™ 8000 Spectrophotometer (ThermoFisher, Waltham, MA, USA). We sequenced RNA using TagSeq, which is a 3’ short transcript method that is a cost-effective alternative to RNAseq and allows for accurate estimates of transcript abundance ^120^. Libraries were prepared for sequencing on one lane of Illumina NovaSeq 6000 SR100 with standard coverage of 3-5 million 100 bp single end reads (University of Texas Austin, Genomic Sequencing and Analysis Facility, Austin, TX, USA). Quality of raw sequences was assessed using *FastQC* ^121^ and visualized with *MultiQC* ^122^. Raw reads were trimmed to remove poly-A tails and adapters and sequences with low quality (phred quality <30) and low complexity (threshold = 50%) using *FastP* ^123^. Reads were then aligned to the *Montipora capitata* reference genome assembly Version 3 ^124^ accessed at (http://cyanophora.rutgers.edu/montipora/; data repository ^114^) and assembled for quantification (exon only assembly) using *HISAT2* ^125^ in combination with *StringTie2* ^126^, and compared to the reference using *GFFcompare* ^127^. Finally, a gene count matrix was generated using *prepDE* ^126^.

#### iv. Metabolomics

Samples for metabolomic analysis were snap-frozen in liquid nitrogen and stored at - 80°C (Fig 1A). Metabolites were extracted from the coral holobiont (homogenized tissue containing host tissue and intracellular symbionts) using 40:40:20 methanol:acetonitrile:water with 0.5%[v/v] formic acid according to ^128^ and identified and quantified at the Rutgers Cancer Institute of New Jersey Metabolomics (New Brunswick, NJ, USA) facility using hydrophilic interaction liquid chromatography separation and liquid chromatography-mass spectrometry (LC-MS) quantification. Metabolite quantification and identification was performed at Rutgers Cancer Institute of New Jersey Metabolomics facility. Hydrophilic interaction liquid chromatography (HILIC) separation was performed on a Vanquish Horizon UHPLC system (Thermo Fisher Scientific, Waltham, MA) with XBridge BEH Amide column (150C:mmC:×C:2.1C:mm, 2.5C:μm particle size, Waters, Milford, MA) using a gradient of solvent A (95%:5% H_2_O:acetonitrile with 20 mM acetic acid, 40 mM ammonium hydroxide, pH 9.4), and solvent B (20%:80% H_2_O:acetonitrile with 20 mM acetic acid, 40 mM ammonium hydroxide, pH 9.4). The gradient was 0 min, 100% B; 3 min, 100% B; 3.2 min, 90% B; 6.2 min, 90% B; 6.5 min, 80% B; 10.5 min, 80% B; 10.7 min, 70% B; 13.5 min, 70% B; 13.7 min, 45% B; 16 min, 45% B; 16.5 min, 100% B and 22 min, 100% B ^129^. The flow rate was 300 μL/min. Injection volume was 5 μL and column temperature was 25 °C. The autosampler temperature was set to 4°C and the injection volume was 5µL. Full scan mass spectrometry analysis was performed on a Thermo Q Exactive PLUS with a HESI source which was set to a spray voltage of −2.7kV under negative mode and 3.5kV under positive mode. The sheath, auxiliary, and sweep gas flow rates of 40, 10, and 2 (arbitrary unit) respectively. The capillary temperature was set to 300°C and aux gas heater was 360°C. The S-lens RF level was 45. The m/z scan range was set to 72 to 1000m/z under both positive and negative ionization mode. The AGC target was set to 3e6, and the maximum IT was 200ms. The resolution was set to 70,000. Full scan data was processed with a targeted data pipeline using MAVEN software package ^130^. The compound identification was assessed using accurate mass and retention time matched to the metabolite standards from the in-house library. Prior to running the samples, the LC-MS system was evaluated for performance readiness by running a commercially available standard mixture and an in-house standard mixture to assess the mass accuracy, signal intensities and the retention time consistency. All known metabolites in the mixture were detected within 5 ppm mass accuracy. Method blank samples matching the composition of the extraction solvent are used in every sample batch to assess background signals and ensure there was no carryover from one run to the next. In addition, the sample queue was randomized with respect to sample treatment to eliminate the potential for batch effects.

#### v. ITS2 symbiont amplicon sequencing

Internal Transcribed Spacer 2 (ITS2) rRNA gene amplicon sequencing was conducted to identify Symbiodiniaceae communities across time points (Fig 1A). A two-step PCR protocol was used for sequencing according to a modification of the Illumina 16S Metagenomic Sequencing Library Preparation protocol (see protocol in OSF data repository ^114^) to amplify the ITS2 region ^131^ of Symbiodiniaceae nuclear ribosomal DNA. A first round of PCR was performed to amplify the ITS2 region of Symbiodiniaceae nuclear ribosomal DNA using ITSintfor2 and ITS2-reverse primers ^132^. Duplicate PCRs were performed on each sample using 4 ng of input DNA in 1uL, 12.5 µL of 1X Phusion HiFi Mastermix (New England Biolabs, Ipswich, MA, Catalog #Cat #M0531S), 0.5 µL each forward and reverse 10 µM primer and 10.5 µL of Ultra-Pure water for a total reaction volume of 25 µL. Negative controls (no template DNA) were performed with Ultra-Pure water instead of DNA. PCR thermocycler protocol included initial denaturation at 95°C for 3 min, 35 cycles of 95°C for 30s, 52°C for 30s, and 72°C for 30s, followed by final extension at 72°C for 2 min. Duplicate PCR products were pooled by sample and run on a 2% agarose gel to confirm successful amplification.

25 µL of each sample was then submitted to the RI-INBRE Molecular Informatics Core at the University of Rhode Island (Kingston, RI, USA) for the index PCR and sequencing. PCR products from the first round of PCR were cleaned with NucleoMag beads (Takara Bio, San Jose, CA, USA) and then visualized by agarose gel electrophoresis. A second round of PCR (50 ng of template DNA, 8 cycles) was performed to attach Nextera XT indices and adapters using the Illumina Nextera XT® Index Kit (Illumina, San Diego, CA) and Phusion HiFi PCR master mix (New England Biolabs, Ipswich, MA, USA, Catalog #Cat #M0531S). PCR products from the second PCR were cleaned with NucleoMag beads and analyzed by agarose gel electrophoresis. Selected samples were confirmed on an Agilent BioAnalyzer DNA1000 chip (Santa Clara, CA, USA). Quantification was performed on all samples prior to pooling using a Qubit fluorometer (Invitrogen, Carlsbad, CA, USA), and the final pooled library was quantified by qPCR in a Roche LightCycler480 with the KAPA Biosystems Illumina Kit (Roche Sequencing, Indianapolis, IN, USA). Samples were sequenced on an Illumina MiSeq platform (Illumina, San Diego, CA, USA) using 2×300 bp paired-end sequencing at the RI-INBRE Molecular Informatics Core (University of Rhode Island, Kingston, RI, USA).

#### vi. 16S rRNA amplicon sequencing

To identify bacterial communities, we conducted 16S rRNA amplicon sequencing across development (Fig 1A). Extracted DNA was prepared using a two-step PCR protocol using published primers 515F (F) and 806RB (R) for the 16S rRNA V4 variable region ^133^. To identify microbial communities, 12 ng per sample of extracted DNA was used in a two-step polymerase chain reaction (PCR) protocol using 515F and 806RB primers for the 16S rRNA V4 variable region ^133^): 515F with Illumina adapter overhang marked in italics (5’-*TCG TCG GCA GCG TCA GAT GTG TAT AAG AGA CAG* GTG CCA GCM GCC GCG GTA A-3’) and 806RB with Illumina adapter overhang marked in italics (5’-*GTC TCG TGG GCT CGG AGA TGT GTA TAA GAG ACA G*GG ACT ACN VGG GTW TCT AAT-3’). Samples were amplified in triplicate (4 ng per replicate) with the following mix for each reaction (25 µL total): 12.5 µL 1X Phusion HiFi Mastermix (New England Biolabs, Ipswich, MA, USA, Catalog #Cat #M0531S), 0.5 µL of each primer (10 µM concentration), 10.5 µL of Ultra-Pure water, and 1 µL of DNA input. All samples were amplified with the following thermocycler conditions: initial denaturation for 2 minutes at 95°C, 35 cycles of 20 seconds at 95°C, 15 seconds at 57°C, and 5 minutes at 72°C, followed by 10 minutes at 72°C. Post-PCR, triplicates were pooled, and amplification was confirmed with a 2% agarose gel. 25 µL of PCR product per sample was then sent for index PCR and sequencing at the RI-INBRE Molecular Informatics Core at the University of Rhode Island (Kingston, RI, USA). PCR products were cleaned with NucleoMag beads (Takara Bio, San Jose, CA, USA) and visualized with agarose gel electrophoresis. 50 ng of cleaned DNA per sample was used in a 8-cycle PCR protocol to attach Nextera XT indices and adapters using the Illumina Nextera XT® Index Kit (Illumina, San Diego, CA, USA, Catalog #FC-131-1024) and Phusion HiFi PCR master mix (New England Biolabs, Ipswich, MA, USA, Catalog #Cat #M0531S). PCR products were then cleaned again with NucleoMag beads and visualized with agarose gel electrophoresis. Quality of selected samples was assessed on an Agilent BioAnalyzer DNA1000 chip (Santa Clara, CA, USA). Quantity was measured for all samples using a Qubit fluorometer (Invitrogen, Carlsbad, CA, USA) prior to pooling. The final pooled library was quantified using qPCR in a Roche LightCycler480 with the KAPA Biosystems Illumina Kit (Roche Sequencing, Indianapolis, IN, USA). Samples were then sequenced on an Illumina MiSeq (Illumina, San Diego, CA, USA) for 2×300 bp paired-end sequencing (University of Rhode Island, Kingston, RI, USA)

### D. Quantification and statistical analyses

#### i. Respirometry

Photosynthesis and respiration rates were extracted using localized linear regressions (LoLinR ^134^) and normalized to average size of each respective life stage to calculate size-normalized respiration and photosynthesis rates (nmol O_2_ mm^-3^ min^-1^). Photosynthesis:respiration ratios (P:R) were calculated as gross photosynthesis divided by respiration. Assumptions of normality and homogeneity of variance for these and following tests were assessed by quantile-quantile plots and Levene’s tests in the *car* package, respectively ^135^. Due to violations in normality and homogeneity of variance, size-normalized respiration and gross photosynthesis rates and P:R ratios were analyzed using Kruskal Wallis rank sum tests in R Statistical Programming v4.2.2 ^136^. Post hoc tests were performed using Dunn tests with Bonferroni P-value adjustment in the *FSA* package ^137^.

#### ii. Size and symbiont density

Size was determined as individual volume by measuring the length and width of the longitudinal and transverse axes of randomly selected individuals from each image (*n* = 15 individuals per time point, *n* = 4 samples per time point) in Fiji/ImageJ ^138,139^. Eggs and embryos (1, 5, and 65 hpf) were approximated as the volume of a sphere, 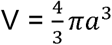, where *a* is the width. Larvae (93-231 hpf) were approximated as a prolate spheroid, 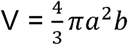, where *a* is 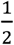 width and *b* is 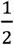 length ^140^. Metamorphosed polyps were approximated as an oblate spheroid, 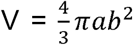, where *a* is 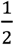 width and *b* is 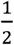 length. We did not statistically compare the size of individual attached recruits on plugs because recruits often settled in aggregation, preventing measurement of individual size. Total surface area of attached recruit tissue on each plug was measured for normalization of symbiont cell density. We tested for differences in size between time points using a non-parametric Kruskal Wallis test. Post hoc tests were performed using Dunn tests with Bonferroni P-value adjustment in the *FSA* package ^137^.

Symbiont cell counts were normalized to the number of individuals in the sample measured by counting all individuals in each sample (egg through metamorphosed polyps; cells individual^-1^). Cell densities were also normalized to the mean size of each life stage (eggs through metamorphosed polyps; cells mm^-3^), or total surface area of tissue sampled (attached recruits; cells mm^-2^). We tested for correlations between size and cells per individual using a Spearman correlation test. Symbiont densities per individual and size-normalized densities were analyzed using a one-way ANOVA test with time point as the main effect, from the stages of eggs to metamorphosed polyps. Post hoc tests were performed using estimated marginal means in the *emmeans* package ^141^. The differences in symbiont cell densities between time points for attached recruits (symbiont cells mm^-2^) were analyzed separately using a non-parametric Kruskal Wallis test.

#### iii. Gene expression in the coral host

Gene expression across development was first visualized using PCA analyses and tested for differences between stages using a permutational analysis of variance (PERMANOVA) in the *vegan* package with variance stabilized transformed gene counts ^142^. Raw gene counts analyzed were analyzed using a likelihood ratio test (LRT) approach to identify differentially expressed genes in the *DESeq2* package ^143^ with a full model design including time point as the main effect and the reduced model removing the effect of time point (reduced=∼1). The LRT approach detects genes that significantly change across the developmental time series by testing whether the main effect of developmental time point (removed in the reduced model) explains a significant amount of variance. Differentially expressed genes (DEGs) were identified as genes with false discovery rate (FDR) adjusted P-value < 0.05.

Patterns of expression of DEGs across development were then visualized using the *DEGreport* package ^144^, which uses hierarchical clustering to identify groups, or clusters, of DEGs that share expression patterns (minimum cluster size = 15). Clusters were assigned to a developmental pattern based on the developmental window during which expression peaked: early development (1-38 hpf), mid development (65-163 hpf) and late development (183-231 hpf). Functional enrichment of genes in each developmental pattern (DEGs peaking in early, mid, or late development) was conducted by analyzing the list of DEGs against all genes expressed and passed filtering using gene ontology (GO) enrichment analysis with length bias adjustment in *goseq* ^145^ and published functional annotation accompanying the *M. capitata* reference genome Version 3 (http://cyanophora.rutgers.edu/montipora/) ^124^; data repository ^114^). Significantly enriched GO terms (Biological Process ontology) were selected as those with false discovery rate adjusted P-value < 0.05. Redundant GO terms were reduced by grouping with semantic similarity (threshold = 0.7, *Caenorhabditis elegans*; ^146^) in the *rrvgo* package ^147^.

#### iv. Metabolomics

Ion counts for metabolites detected at both positive and negative polarities in each sample were center normalized to the respective sample median ion count to account for variability in sample concentrations (*n* = 183 total detected and annotated metabolites). For metabolites that were detected at both positive and negative polarities, we selected the normalized ion counts from the polarity with the strongest detection (i.e., greatest normalized ion count) for each metabolite. Following normalization and polarity selection, normalized ion counts were then log_10_ transformed. We examined holobiont metabolomic profiles across development by first examining unsupervised PCA visualizations and PERMANOVA tests in the vegan package ^142^. Metabolomic profiles were then analyzed using supervised partial least squares discriminant analysis (PLS-DA) followed by identification of metabolite variables of importance in projection (VIP) in the mixOmics ^148^ and RVAideMemoire ^149^ packages. The VIP score (calculated as the sum of squared correlations between the PLS-DA components and the individual variable) indicates a metabolite’s importance in discriminating between time points in the PLS-DA model. Metabolites with a VIP score greater than or equal to 1 (VIP ≥ 1) are statistically important in discriminating variation due to developmental time point ^150^. The list of significant VIP metabolites was then analyzed for compound class and KEGG pathway enrichment in the MetaboAnalyst 5.0 web interface ^151^. The effect of time point on pool size of specific VIP metabolites of interest (including glucose, glutamine, glutamate, and arginine-glutamine) was tested using one-way ANOVA tests with post hoc comparisons conducted using estimated marginal means tests in the emmeans package in R ^141^.

#### v. ITS2 symbiont amplicon sequencing

Sequences were quality controlled and subsampled to even sequencing depth of 10,551 (the minimum sequencing depth in our samples) and analyzed with the SymPortal workflow ^34^ locally to identify Symbiodiniaceae Defining Intragenomic Variants (DIVs) that determine ITS2 type profiles. Relative abundance was calculated and visualized for DIVs at >1% relative abundance. Alpha diversity of DIVs was calculated using Shannon and Inverse Simpson indices and analyzed using one-way ANOVA with time point as the main effect. Shifts in Symbiodiniaceae communities (based on Bray-Curtis dissimilarity) were visualized across ontogeny using a non-metric multidimensional scaling analysis (NMDS) in the *phyloseq* package ^152^. Changes in the relative abundance of major ITS2 profiles and the ratio of major profiles were analyzed using one-way ANOVA with time point as the main effect. Post hoc tests were conducted in the *emmeans* package ^141^.

#### vi. 16S rRNA amplicon sequencing

We conducted data analysis using the *mothur* analytical pipeline (v1.46.1) ^153^. Contigs were assembled using trimoverlap=TRUE and primers were removed. Contigs were then filtered to remove sequences with ambiguous calls and sequences <200 bp and >350 bp length. Unique sequences were identified and aligned to the V4 region of the Silva database (release 138.1 ^154^) using the *mothur*-formatted version of the RDP training set (v.9; https://mothur.org/wiki/rdp_reference_files/). Sequences that did not align within the region of interest and those with >8 polymers (maxhomop=8) were removed. Sequences were then pre-clustered with nucleotide differences of 1 (diffs=1). Chimeras were removed using VSEARCH ^155^ in *mothur* (https://github.com/torognes/vsearch). Remaining sequences were then classified to the Silva database (release 138.1) using the *mothur*-formatted version of the RDP training set (v.9) removing sequences classified as chloroplast, mitochondria, Archaea, Eukaryote, or unknown domain. 13% of sequences passed these QC and filtering parameters due to a high proportion of sequences that did not align to the Silva database, which may have been due to unintentional amplification of host sequences. Remaining sequences were clustered to operational taxonomic units (OTU; cutoff=0.03) and subsampled to 900 sequences to obtain even sampling depth. Due to low bacterial sequence reads, 13 out of 39 samples were removed after subsampling. The removed samples included egg (1 hpf), embryo (5 hpf), and half of the samples at 38 and 65 hpf stages, indicating low bacterial sequences generated from the earliest life stages in the time series, which corresponded to samples with low DNA concentration during extraction. These samples were not included in further analysis. Bacterial OTU counts were analyzed in R using the *phyloseq* package ^152^. Shifts in bacterial communities (calculated using Bray-Curtis dissimilarity) across ontogeny were visualized using a non-metric multidimensional scaling analysis (NMDS) and analyzed using PERMANOVA tests. Taxa abundance was calculated as percent relative abundance and filtered to those present at >0.5% for visualization at the genus level. Alpha diversity was calculated using Shannon and Inverse Simpson indices. Changes in alpha diversity were analyzed using a one-way ANOVA with time point as the main effect.

## Supplemental information titles and legends

**Fig S1. Symbiont cell density per individual from egg to metamorphosed polyp stages.** Data are represented as mean ± standard error. Life stage is indicated on the x-axis as hours post fertilization (hpf). Color corresponds to major life history grouping (eggs=brown, embryos=yellow, larvae=cyan, metamorphosed polyps=green, attached recruits=pink).

**Fig S2. Symbiodiniaceae ITS2 analysis.** (A) NMDS plot of Symbiodiniaceae communities across development (eggs=brown, embryos=yellow, larvae=cyan, metamorphosed polyps=green, attached recruits=pink). Shape indicates life stage. (B) Heatmap of ITS2 DIV relative abundance of taxa abundant at >1% relative abundance across life stages. Red = relative abundance of 1; Blue = relative abundance of 0. (C) Ratio of *Cladocopium* sp. to *Durusdinium glynii* relative abundance across life stages. Error bars indicate standard error of mean. Shared letters indicate groups are not significantly different. Groups with red letters are significantly different at post hoc p<0.05.

**Fig S3. Bacterial community 16S analysis.** (A) NMDS plot of bacterial communities across development (eggs=brown, embryos=yellow, larvae=cyan, metamorphosed polyps=green, attached recruits=pink). Shape indicates life stage. (B) Alpha diversity metrics of bacterial communities calculated as Shannon (top) and Inverse Simpson (bottom) metrics. (C) Heatmap of bacterial relative abundance across life stages at the phylum level. (D) Heatmap of bacterial relative abundance across life stages at the family:genus level in taxa at greater than 0.5% relative abundance. Red colors indicate higher relative abundance.

**Fig S4. Differentially expressed genes across development.** Heatmap of expression (z-score) of differentially expressed genes (9,181 total DEGs; y-axis) across time points (x-axis) identified by likelihood ratio tests (FDR P-value < 0.05). Within the heatmap, orange indicates increased expression with purple indicating reduced expression relative to the average of that gene’s expression in all samples. Column annotation colors indicate life stages.

**Fig S5. Expression patterns of gene clusters across development.** Panels show detected clusters (minimum number of genes = 15) of genes that share expression patterns across life stages. Expression displayed as z-score of variance stabilized transformed gene counts. Dotted lines indicate 38 hpf and 163 hpf that separate early, mid, and late developmental stages. Clusters were assigned to developmental patterns based on the time at which expression peaked: early development (1-38 hpf), mid development (65-163 hpf) and late development (183-231 hpf). Red lines indicate expression for individual genes with black lines indicating the cluster mean expression across development.

**Fig S6. Functional enrichment of genes in early development.** Lollipop plot of functional enrichment of genes peaking in early development (1-38 hpf). Plot shows significant (FDR P<0.05) gene ontology (GO) terms (Biological Process; BP). x-axis and dot size indicate % hits (percentage of the number of DEGs in the respective GO category compared to total number of genes included in that category) with color indicating P-value. Parent term BP is indicated in panels on the right and individual GO terms on the left.

**Fig S7. Metabolite variables of importance in project (VIP) scores for all metabolites.** Red dashed line indicates VIP = 1, above which metabolites are statistically important in discriminating between life stage groups. VIP scores calculated for components 1 and 2 in the partial least squares discriminant model.

**Table S1. Sample size of response variables across developmental stages.** Time point identified by life stage group and hours post-fertilization (hpf). Shaded boxes indicate that the response was not measured at the respective time point.

**Table S2. Gene expression patterns across development.** Number of genes identified in each gene expression cluster and the assignment of each cluster to developmental patterns. Developmental patterns describe clusters of genes that peaked in expression in early development (1-38 hpf), mid development (65-163 hpf), and late development (183-231 hpf) periods.

**Table S3. Functional enrichment of genes in early development.** Significantly enriched (adjusted P-value < 0.05) Biological Process (BP) gene ontology (GO) terms in *Montipora capitata* early developmental stages (1-38 hpf). Functional enrichment analyses performed in the goseq package in R with redundant terms reduced by grouping with semantic similarity in the rrvgo package in R to generate parent terms. Percent Hits calculated as the number of differentially expressed genes in the GO category divided by the total number of genes from the background set in the GO category.

**Table S4. Functional enrichment of genes in mid-development.** Significantly enriched (adjusted P-value < 0.05) Biological Process (BP) gene ontology (GO) terms in *Montipora capitata* mid-developmental stages (65-163 hpf). Functional enrichment analyses performed in the goseq package in R with redundant terms reduced by grouping with semantic similarity in the rrvgo package in R to generate parent terms. Percent Hits calculated as the number of differentially expressed genes in the GO category divided by the total number of genes from the background set in the GO category.

**Table S5. Functional enrichment of genes in late development.** Significantly enriched (adjusted P-value < 0.05) Biological Process (BP) gene ontology (GO) terms in *Montipora capitata* late developmental stages (183-231 hpf). Functional enrichment analyses performed in the goseq package in R with redundant terms reduced by grouping with semantic similarity in the rrvgo package in R to generate parent terms. Percent Hits calculated as the number of differentially expressed genes in the GO category divided by the total number of genes from the background set in the GO category.

**Table S6. Metabolites that significantly vary across development.** Variable importance in projection (VIP) values for metabolites that significantly (VIP ≥ 1) contribute to discrimination between life stages as determined by partial least squares discriminant analyses. Pathway or function indicates metabolic function of each metabolite. Ontogenetic peak indicates developmental periods in which metabolite pool size was highest.

**Table S7. Class enrichment of metabolites that vary across development.** Enrichment of metabolite compound classes (sub-class) of VIP metabolites conducted using Metaboanalyst v5.0 web interface. Bold indicates classes significantly enriched at FDR p-value < 0.05.

**Table S8. KEGG enrichment of metabolites that vary across development.** Enrichment of KEGG pathways of VIP metabolites conducted using Metaboanalyst v5.0 web interface. Bold indicates pathways significantly enriched at FDR p-value < 0.05.

## Notes

https://github.com/AHuffmyer/EarlyLifeHistory_Energetics/releases/tag/v2.1.1

https://doi.org/10.17605/OSF.IO/KMU7H

https://www.ncbi.nlm.nih.gov/bioproject/PRJNA900235/

### Competing Interest Statement

The authors have declared no competing interest.

### Summary of Updates

This manuscript has been revised to include differential gene expression analysis, supervised linear discriminant metabolomics analysis, and revisions of main and supplemental figures. In this revision, the results and discussion sections have been revised to provide updated discussion of findings and the methods section now follows the STAR methods format.

## References

1. Muscatine, L., and Porter, J.W. (1977). Reef Corals: Mutualistic Symbioses Adapted to Nutrient-Poor Environments. Bioscience 27, 454–460. 10.2307/1297526.

2. Yellowlees, D., Rees, T.A.V., and Leggat, W. (2008). Metabolic interactions between algal symbionts and invertebrate hosts. Plant Cell Environ. 31, 679–694. 10.1111/j.1365-3040.2008.01802.x.

3. van der Heijden, M.G.A., Dombrowski, N., and Schlaeppi, K. (2017). Continuum of root-fungal symbioses for plant nutrition. Proc. Natl. Acad. Sci. U. S. A. 114, 11574–11576. 10.1073/pnas.1716329114.

4. Dubilier, N., Bergin, C., and Lott, C. (2008). Symbiotic diversity in marine animals: the art of harnessing chemosynthesis. Nat. Rev. Microbiol. 6, 725–740. 10.1038/nrmicro1992

5. LaJeunesse, T.C., Parkinson, J.E., Gabrielson, P.W., Jeong, H.J., Reimer, J.D., Voolstra, C.R., and Santos, S.R. (2018). Systematic revision of Symbiodiniaceae highlights the antiquity and diversity of coral endosymbionts. Curr. Biol. 28, 2570–2580. 10.1016/j.cub.2018.07.008.

6. Muscatine, L., Mccloskey, L.R., and Marian, R.E. (1981). Estimating the daily contribution of carbon from zooxanthellae to coral animal respiration. Limnol. Oceanogr. 26, 601–611. 10.4319/lo.1981.26.4.0601.

7. Falkowski, P.G., Dubinsky, Z., Muscatine, L., and Porter, J.W. (2006). Light and the bioenergetics of a symbiotic coral. Bioscience 34, 705–709. 10.2307/1309663.

8. Falkowski, P.G., Dubinsky, Z., Muscatine, L., and McCloskey, L. (1993). Population control in symbiotic corals - Ammonium ions and organic materials maintain the density of zooxanthellae. Bioscience 43, 606–611. 10.2307/1312147.

9. Roberty, S., Béraud, E., Grover, R., and Ferrier-Pagès, C. (2020). Coral Productivity Is Co-Limited by Bicarbonate and Ammonium Availability. Microorganisms 8, 640. 10.3390/microorganisms8050640.

10. Bourne, D.G., Morrow, K.M., and Webster, N.S. (2016). Insights into the Coral Microbiome: Underpinning the Health and Resilience of Reef Ecosystems. Annu. Rev. Microbiol. 70, 317–340. 10.1146/annurev-micro-102215-095440.

11. Connolly, S.R., Bellwood, D.R., and Hughes, T.P. (2003). Indo-pacific biodiversity of coral reefs: Deviations from a mid-domain model. Ecology 84, 2178–2190. 10.1890/02-0254.

12. Woodhead, A.J., Hicks, C.C., Norström, A.V., Williams, G.J., and Graham, N.A.J. (2019). Coral reef ecosystem services in the Anthropocene. Funct. Ecol. 33, 1023–1034. 10.1111/1365-2435.13331.

13. Weis, V.M. (2008). Cellular mechanisms of Cnidarian bleaching: stress causes the collapse of symbiosis. J. Exp. Biol. 211, 3059–3066. 10.1242/jeb.009597.

14. Sully, S., Burkepile, D.E., Donovan, M.K., Hodgson, G., and van Woesik, R. (2019). A global analysis of coral bleaching over the past two decades. Nat. Commun. 10, 1264. 10.1038/s41467-019-09238-2.

15. Eddy, T.D., Lam, V.W.Y., Reygondeau, G., Cisneros-Montemayor, A.M., Greer, K., Palomares, M.L.D., Bruno, J.F., Ota, Y., and Cheung, W.W.L. (2021). Global decline in capacity of coral reefs to provide ecosystem services. One Earth 4, 1278–1285. 10.1016/j.oneear.2021.08.016.

16. Ritson-Williams, R., Arnold, S.N., Fogarty, N.D., Steneck, R.S., Vermeij, M.J.A., and Paul, V.J. (2009). New perspectives on ecological mechanisms affecting coral recruitment on reefs. Smithson. Contrib. Mar. Sci. 38, 437–457. 10.5479/si.01960768.38.437.

17. Pineda, J. (2000). Linking larval settlement to larval transport: assumptions, potentials, and pitfalls. Oceanography of the eastern Pacific 1, 84–105.

18. Marshall, D.J., Pettersen, A.K., Bode, M., and White, C.R. (2020). Developmental cost theory predicts thermal environment and vulnerability to global warming. Nat. Ecol. Evol. 4, 406–411. 10.1038/s41559-020-1114-9.

19. Drake, J.L., Mass, T., Stolarski, J., Von Euw, S., van de Schootbrugge, B., and Falkowski, P.G. (2020). How corals made rocks through the ages. Glob. Chang. Biol. 26, 31–53. 10.1111/gcb.14912.

20. Baird, A.H., Guest, J.R., and Willis, B.L. (2009). Systematic and Biogeographical Patterns in the Reproductive Biology of Scleractinian Corals. Annu. Rev. Ecol. Evol. Syst. 40, 551–571. 10.1146/annurev.ecolsys.110308.120220.

21. Rosset, S.L., Oakley, C.A., Ferrier-Pagès, C., Suggett, D.J., Weis, V.M., and Davy, S.K. (2020). The Molecular Language of the Cnidarian–Dinoflagellate Symbiosis. Trends Microbiol. 29, 320–333. 10.1016/j.tim.2020.08.005.

22. Edmunds, P.J. (2023). Coral recruitment: patterns and processes determining the dynamics of coral populations. Biol. Rev. Camb. Philos. Soc. 98, 1862–1886. 10.1111/brv.12987.

23. Harii, S., Nadaoka, K., Yamamoto, M., and Iwao, K. (2007). Temporal changes in settlement, lipid content and lipid composition of larvae of the spawning hermatypic coral Acropora tenuis. Mar. Ecol. Prog. Ser. 346, 89–96. 10.3354/meps07114.

24. Rivest, E.B., Chen, C.-S., Fan, T.-Y., Li, H.-H., and Hofmann, G.E. (2017). Lipid consumption in coral larvae differs among sites: a consideration of environmental history in a global ocean change scenario. Proc. R. Soc. Lond. Ser. B. Biol. Sci. 284, 20162825. 10.1098/rspb.2016.2825.

25. Graham, E.M., Baird, A.H., Connolly, S.R., Sewell, M.A., and Willis, B.L. (2016). Uncoupling temperature-dependent mortality from lipid depletion for scleractinian coral larvae. Coral Reefs 36, 97–104. 10.1007/s00338-016-1501-5.

26. Figueiredo, J., Baird, A.H., Cohen, M.F., Flot, J.-F., Kamiki, T., Meziane, T., Tsuchiya, M., and Yamasaki, H. (2012). Ontogenetic change in the lipid and fatty acid composition of scleractinian coral larvae. Coral Reefs 31, 613–619. 10.1007/s00338-012-0874-3.

27. Gaither, M.R., and Rowan, R. (2010). Zooxanthellar symbiosis in planula larvae of the coral *Pocillopora damicornis*. J. Exp. Mar. Bio. Ecol. 386, 45–53. 10.1016/j.jembe.2010.02.003.

28. Harii, S., Yamamoto, M., and Hoegh-Guldberg, O. (2010). The relative contribution of dinoflagellate photosynthesis and stored lipids to the survivorship of symbiotic larvae of the reef-building corals. Mar. Biol. 157, 1215–1224. 10.1007/s00227-010-1401-0.

29. Hazraty-Kari, S., Masaya, M., Kawachi, M., and Harii, S. (2022). The early acquisition of symbiotic algae benefits larval survival and juvenile growth in the coral *Acropora tenuis*. J. Exp. Zool. A. Ecol. Integr. Physiol. 337, 559–565. 10.1002/jez.2589.

30. Kopp, C., Domart-Coulon, I., Barthelemy, D., and Meibom, A. (2016). Nutritional input from dinoflagellate symbionts in reef-building corals is minimal during planula larval life stage. Sci Adv 2, e1500681. 10.1126/sciadv.1500681.

31. Huffmyer, A., Ashey, J., Strand, E., Chiles, E., Su, X., and Putnam, H. (2024). Coral larvae increase nitrogen assimilation to stabilize algal symbiosis and combat bleaching under increased temperature. PLoS Biol. 22, e3002875. 10.1371/journal.pbio.3002875.

32. LaJeunesse, T.C., and Thornhill, D.J. (2011). Improved resolution of reef-coral endosymbiont (Symbiodinium) species diversity, ecology, and evolution through psbA non-coding region genotyping. PLoS One 6, e29013. 10.1371/journal.pone.0029013.

33. Wham, D.C., Ning, G., and LaJeunesse, T.C. (2017). *Symbiodinium glynnii* sp. nov., a species of stress-tolerant symbiotic dinoflagellates from pocilloporid and montiporid corals in the Pacific Ocean. Phycologia 56, 396–409. 10.2216/16-86.1.

34. Hume, B.C.C., Smith, E.G., Ziegler, M., Warrington, H.J.M., Burt, J.A., LaJeunesse, T.C., Wiedenmann, J., and Voolstra, C.R. (2019). SymPortal: A novel analytical framework and platform for coral algal symbiont next-generation sequencing ITS2 profiling. Mol. Ecol. Resour. 19, 1063–1080. 10.1111/1755-0998.13004.

35. Burriesci, M.S., Raab, T.K., and Pringle, J.R. (2012). Evidence that glucose is the major transferred metabolite in dinoflagellate-cnidarian symbiosis. J. Exp. Biol. 215, 3467–3477. 10.1242/jeb.070946.

36. Gordon, B.R., and Leggat, W. (2010). Symbiodinium—Invertebrate Symbioses and the Role of Metabolomics. Mar. Drugs 8, 2546–2568. 10.3390/md810546.

37. Hillyer, K.E., Dias, D., Lutz, A., Roessner, U., and Davy, S.K. (2018). 13C metabolomics reveals widespread change in carbon fate during coral bleaching. Metabolomics 14, 12. 10.1007/s11306-017-1306-8

38. Pernice, M., Meibom, A., Van Den Heuvel, A., Kopp, C., Domart-Coulon, I., Hoegh-Guldberg, O., and Dove, S. (2012). A single-cell view of ammonium assimilation in coral– dinoflagellate symbiosis. ISME J. 6, 1314–1324. 10.1038/ismej.2011.196.

39. Rädecker, N., Pogoreutz, C., Gegner, H.M., Cárdenas, A., Roth, F., Bougoure, J., Guagliardo, P., Wild, C., Pernice, M., Raina, J.-B., et al. (2021). Heat stress destabilizes symbiotic nutrient cycling in corals. Proc. Natl. Acad. Sci. U. S. A. 118, e2022653118. 10.1073/pnas.2022653118.

40. Chiles, E.N., Huffmyer, A.S., Drury, C., Putnam, H.M., Bhattacharya, D., and Su, X. (2022). Stable isotope tracing reveals compartmentalized nitrogen assimilation in scleractinian corals. Front. Mar. Sci., 9, 1035523. 10.3389/fmars.2022.1035523

41. Padilla-Gamiño, J.L., Bidigare, R.R., Barshis, D.J., Alamaru, A., Hédouin, L., Hernández-Pech, X., Kandel, F., Leon Soon, S., Roth, M.S., Rodrigues, L.J., et al. (2013). Are all eggs created equal? A case study from the Hawaiian reef-building coral *Montipora capitata*. Coral Reefs 32, 137–152. 10.1007/s00338-012-0957-1.

42. Buttari, F., Tsai, S., Wen, Z.-H., Cheng, J.-O., and Lin, C. (2024). Lipid profiling during embryogenesis of coral *Galaxea fascicularis*. Mar. Biol. 171, 199. 10.1007/s00227-024-04521-3.

43. Okubo, N., Mezaki, T., Nozawa, Y., Nakano, Y., Lien, Y.-T., Fukami, H., Hayward, D.C., and Ball, E.E. (2013). Comparative embryology of eleven species of stony corals (Scleractinia). PLoS One 8, e84115. 10.1371/journal.pone.0084115.

44. Reyes-Bermudez, A., Villar-Briones, A., Ramirez-Portilla, C., Hidaka, M., and Mikheyev, A.S. (2016). Developmental Progression in the Coral *Acropora digitifera* Is Controlled by Differential Expression of Distinct Regulatory Gene Networks. Genome Biol. Evol. 8, 851– 870. 10.1093/gbe/evw042.

45. Heather, Q.M., and Mark, Q.M. (2007). Embryonic development in two species of scleractinian coral embryos: *Symbiodinium* localization and mode of gastrulation. Evol. Dev. 9, 355–367. 10.1111/j.1525-142X.2007.00173.x.

46. Fadlallah, Y.H. (1983). Coral Reefs Sexual Reproduction , Development and Larval Biology in Scleractinian Corals A Review. Coral Reefs 2, 129–150. 10.1007/BF00336720.

47. Ghosh, S., Körte, A., Serafini, G., Yadav, V., and Rodenfels, J. (2022). Developmental energetics: Energy expenditure, budgets and metabolism during animal embryogenesis. Semin. Cell Dev. Biol. 138, 83–93. 10.1016/j.semcdb.2022.03.009.

48. Rodrigues, L.J., and Padilla-Gamiño, J.L. (2022). Trophic provisioning and parental trade-offs lead to successful reproductive performance in corals after a bleaching event. Sci. Rep. 12, 18702. 10.1038/s41598-022-21998-4.

49. Graham, E.M., Baird, A.H., Connolly, S.R., Sewell, M.A., and Willis, B.L. (2013). Rapid declines in metabolism explain extended coral larval longevity. Coral Reefs 32, 539–549. 10.1007/s00338-012-0999-4.

50. Christiansen, F.B., and Fenchel, T.M. (1979). Evolution of marine invertebrate reproductive patterns. Theor. Popul. Biol. 16, 267–282. 10.1016/0040-5809(79)90017-0.

51. Hadfield, M., and Strathmann, M. (1996). Variability, flexibility and plasticity in life histories of marine invertebrates. Oceanol. Acta 19, 323–334.

52. Axworthy, J.B., Timmins-Schiffman, E., Brown, T., Rodrigues, L.J., Nunn, B.L., and Padilla-Gamiño, J.L. (2022). Shotgun Proteomics Identifies Active Metabolic Pathways in Bleached Coral Tissue and Intraskeletal Compartments. Front. Mar. Sci. 9, 797517.

53. DeSalvo, M.K., Sunagawa, S., Voolstra, C.R., and Medina, M. (2010). Transcriptomic responses to heat stress and bleaching in the elkhorn coral *Acropora palmata*. Mar. Ecol. Prog. Ser. 402, 97–113.

54. Kenkel, C.D., Meyer, E., and Matz, M.V. (2013). Gene expression under chronic heat stress in populations of the mustard hill coral (*Porites astreoides*) from different thermal environments. Mol. Ecol. 22, 4322–4334. 10.1111/mec.12390.

55. Petrou, K., Nunn, B.L., Padula, M.P., Miller, D.J., and Nielsen, D.A. (2021). Broad scale proteomic analysis of heat-destabilised symbiosis in the hard coral *Acropora millepora*. Sci. Rep. 11, 19061. 10.1038/s41598-021-98548-x.

56. Leinbach, S.E., Speare, K.E., Rossin, A.M., Holstein, D.M., and Strader, M.E. (2021). Energetic and reproductive costs of coral recovery in divergent bleaching responses. Sci. Rep. 11, 23546. 10.1038/s41598-021-02807-w.

57. Radecker, N., Pogoreutz, C., Voolstra, C.R., Wiedenmann, J., and Wild, C. (2015). Nitrogen cycling in corals: The key to understanding holobiont functioning? Trends Microbiol. 23, 490–497. 10.1016/j.tim.2015.03.008.

58. Rädecker, N., Escrig, S., Spangenberg, J.E., Voolstra, C.R., and Meibom, A. (2023). Coupled carbon and nitrogen cycling regulates the cnidarian–algal symbiosis. Nat. Commun. 14, 6948. 10.1038/s41467-023-42579-7.

59. Cui, G., Mi, J., Moret, A., Menzies, J., Zhong, H., Li, A., Hung, S.-H., Al-Babili, S., and Aranda, M. (2023). A carbon-nitrogen negative feedback loop underlies the repeated evolution of cnidarian–Symbiodiniaceae symbioses. Nat. Commun. 14, 6949. 10.1038/s41467-023-42582-y.

60. Cui, G., Liew, Y.J., Li, Y., Kharbatia, N., Zahran, N.I., Emwas, A.-H., Eguiluz, V.M., and Aranda, M. (2019). Host-dependent nitrogen recycling as a mechanism of symbiont control in Aiptasia. PLoS Genet. 15, e1008189. 10.1371/journal.pgen.1008189.

61. Rahav, O., Dubinsky, Z., Achituv, Y., Falkowski, P.G., and Smith, D.C. (1989). Ammonium metabolism in the zooxanthellate coral, *Stylophora Pistillata*. Proc. R. Soc. Lond. Ser. B, Biol. Sci. 236, 325–337. 10.1098/rspb.1989.0026.

62. Morris, L.A., Voolstra, C.R., Quigley, K.M., Bourne, D.G., and Bay, L.K. (2019). Nutrient Availability and Metabolism Affect the Stability of Coral–Symbiodiniaceae Symbioses. Trends Microbiol. 27, 678–689. 10.1016/j.tim.2019.03.004.

63. Roberts, J.M., Fixter, L.M., and Davies, P.S. (2001). Ammonium metabolism in the symbiotic sea anemone *Anemonia viridis*. Hydrobiologia 461, 25–35. 10.1023/a:1012752828587.

64. Su, Y., Zhou, Z., and Yu, X. (2018). Possible roles of glutamine synthetase in responding to environmental changes in a scleractinian coral. Mol. Biol. Rep. 45, 2115–2124. 10.1007/s11033-018-4369-3.

65. Cui, G., Liew, Y.J., Konciute, M.K., Zhan, Y., Hung, S.-H., Thistle, J., Gastoldi, L., Schmidt-Roach, S., Dekker, J., and Aranda, M. (2022). Nutritional control regulates symbiont proliferation and life history in coral-dinoflagellate symbiosis. BMC Biol. 20, 103. 10.1186/s12915-022-01306-2.

66. Krueger, T., Horwitz, N., Bodin, J., Giovani, M.-E., Escrig, S., Fine, M., and Meibom, A. (2020). Intracellular competition for nitrogen controls dinoflagellate population density in corals. Proc. Biol. Sci. 287, 20200049. 10.1098/rspb.2020.0049.

67. Xiang, T., Lehnert, E., Jinkerson, R.E., Clowez, S., Kim, R.G., DeNofrio, J.C., Pringle, J.R., and Grossman, A.R. (2020). Symbiont population control by host-symbiont metabolic interaction in Symbiodiniaceae-cnidarian associations. Nat. Commun. 11, 108. 10.1038/s41467-019-13963-z.

68. Zoccola, D., Ganot, P., Bertucci, A., Caminiti-Segonds, N., Techer, N., Voolstra, C.R., Aranda, M., Tambutté, E., Allemand, D., Casey, J.R., et al. (2015). Bicarbonate transporters in corals point towards a key step in the evolution of cnidarian calcification. Sci. Rep. 5, 9983. 10.1038/srep09983.

69. Tambutté, S., Holcomb, M., Ferrier-Pagès, C., Reynaud, S., Tambutté, É., Zoccola, D., and Allemand, D. (2011). Coral biomineralization: From the gene to the environment. J. Exp. Mar. Bio. Ecol. 408, 58–78. 10.1016/j.jembe.2011.07.026.

70. Levy, S., Elek, A., Grau-Bové, X., Menéndez-Bravo, S., Iglesias, M., Tanay, A., Mass, T., and Sebé-Pedrós, A. (2021). A stony coral cell atlas illuminates the molecular and cellular basis of coral symbiosis, calcification, and immunity. Cell 184, 2973–2987.e18. 10.1016/j.cell.2021.04.005.

71. Matthews, J.L., Oakley, C.A., Lutz, A., Hillyer, K.E., Roessner, U., Grossman, A.R., Weis, V.M., and Davy, S.K. (2018). Partner switching and metabolic flux in a model cnidarian-dinoflagellate symbiosis. Proc. Biol. Sci. 285, 20182336. 10.1098/rspb.2018.2336.

72. Fürst, P. (2001). New developments in glutamine delivery. J. Nutr. 131, 2562S – 8S. 10.1093/jn/131.9.2562S.

73. Jaffe, M.D., Padilla-Gamiño, J.L., Nunn, B.L., and Rodrigues, L.J. (2023). Coral trophic pathways impact the allocation of carbon and nitrogen for egg development after bleaching. Front. Ecol. Evol. 11, 1251220. 10.3389/fevo.2023.1251220.

74. Cunning, R., Ritson-Williams, R., and Gates, R.D. (2016). Patterns of bleaching and recovery of *Montipora capitata* in Kane’ohe Bay, Hawai’i, USA. Mar. Ecol. Prog. Ser. 551, 131–139. 10.3354/meps11733.

75. Wall, C.B., Kaluhiokalani, M., Popp, B.N., Donahue, M.J., and Gates, R.D. (2020). Divergent symbiont communities determine the physiology and nutrition of a reef coral across a light-availability gradient. ISME J. 14, 945–958. 10.1038/s41396-019-0570-1.

76. Roach, T.N.F., Dilworth, J., Christian Martin, H., Daniel Jones, A., Quinn, R., and Drury, C. (2021). Metabolomic signatures of coral bleaching history. Nat. Ecol. Evol. 5, 495–503. 10.1101/2020.05.10.087072.

77. Dilworth, J., Caruso, C., Kahkejian, V.A., Baker, A.C., and Drury, C. (2021). Host genotype and stable differences in algal symbiont communities explain patterns of thermal stress response of *Montipora capitata* following thermal pre-exposure and across multiple bleaching events. Coral Reefs 40, 151–163. 10.1007/s00338-020-02024-3.

78. Padilla-Gamiño, J.L., Pochon, X., Bird, C., Concepcion, G.T., and Gates, R.D. (2012). From parent to gamete: vertical transmission of *Symbiodinium* (Dinophyceae) ITS2 sequence assemblages in the reef building coral *Montipora capitata*. PLoS One 7, e38440. 10.1371/journal.pone.0038440.

79. Ng, T.Y., Chui, A.P.Y., Tsang, R.H.L., Yeung, C.W., and Ang, P. (2024). Hong Kong coral recruits do not gain advantage by being flexible in establishing symbiosis under simulated climate change conditions. J. Coast. Res. 113, 936–940. 10.2112/jcr-si113-183.1.

80. Allen-Waller, L., and Barott, K.L. (2023). Symbiotic dinoflagellates divert energy away from mutualism during coral bleaching recovery. Symbiosis 89, 173–186. 10.1007/s13199-023-00901-3.

81. Glaze, T.D., Erler, D.V., and Siljanen, H.M.P. (2022). Microbially facilitated nitrogen cycling in tropical corals. ISME J. 16, 68–77. 10.1038/s41396-021-01038-1.

82. Peixoto, R.S., Rosado, P.M., Leite, D.C. de A., Rosado, A.S., and Bourne, D.G. (2017). Beneficial Microorganisms for Corals (BMC): Proposed Mechanisms for Coral Health and Resilience. Front. Microbiol. 8, 341. 10.3389/fmicb.2017.00341.

83. Reich, H.G., Camp, E.F., Roger, L.M., and Putnam, H.M. (2022). The trace metal economy of the coral holobiont: supplies, demands and exchanges. Biol. Rev. Camb. Philos. Soc. 98, 623–642. 10.1111/brv.12922.

84. Ceh, J., Kilburn, M.R., Cliff, J.B., Raina, J.B., Van Keulen, M., and Bourne, D.G. (2013). Nutrient cycling in early coral life stages: *Pocillopora damicornis* larvae provide their algal symbiont (*Symbiodinium*) with nitrogen acquired from bacterial associates. Ecol. Evol. 3, 2393–2400. 10.1002/ece3.642.

85. Matthews, J.L., Raina, J.-B., Kahlke, T., Seymour, J.R., van Oppen, M.J.H., and Suggett, D.J. (2020). Symbiodiniaceae-bacteria interactions: rethinking metabolite exchange in reef-building corals as multi-partner metabolic networks. Environ. Microbiol. 22, 1675–1687. 10.1111/1462-2920.14918.

86. Kimes, N.E., Van Nostrand, J.D., Weil, E., Zhou, J., and Morris, P.J. (2010). Microbial functional structure of *Montastraea faveolata*, an important Caribbean reef-building coral, differs between healthy and yellow-band diseased colonies. Environ. Microbiol. 12, 541–556. 10.1111/j.1462-2920.2009.02113.x.

87. Raina, J.-B., Dinsdale, E.A., Willis, B.L., and Bourne, D.G. (2010). Do the organic sulfur compounds DMSP and DMS drive coral microbial associations? Trends Microbiol. 18, 101–108. 10.1016/j.tim.2009.12.002.

88. Raina, J.-B., Tapiolas, D., Motti, C.A., Foret, S., Seemann, T., Tebben, J., Willis, B.L., and Bourne, D.G. (2016). Isolation of an antimicrobial compound produced by bacteria associated with reef-building corals. PeerJ 4, e2275. 10.7717/peerj.2275.

89. Matthews, J.L., Khalil, A., Siboni, N., Bougoure, J., Guagliardo, P., Kuzhiumparambil, U., DeMaere, M., Le Reun, N.M., Seymour, J.R., Suggett, D.J., et al. (2023). Coral endosymbiont growth is enhanced by metabolic interactions with bacteria. Nat. Commun. 14, 6864. 10.1038/s41467-023-42663-y.

90. Cárdenas, A., Raina, J.-B., Pogoreutz, C., Rädecker, N., Bougoure, J., Guagliardo, P., Pernice, M., and Voolstra, C.R. (2022). Greater functional diversity and redundancy of coral endolithic microbiomes align with lower coral bleaching susceptibility. ISME J. 16, 2406– 2420. 10.1038/s41396-022-01283-y.

91. Sogin, E.M., Putnam, H.M., Nelson, C.E., Anderson, P., and Gates, R.D. (2017). Correspondence of coral holobiont metabolome with symbiotic bacteria, archaea and Symbiodinium communities. Environ. Microbiol. Rep. 9, 310–315. 10.1111/1758-2229.12541.

92. Engelberts, J.P., Robbins, S.J., Damjanovic, K., and Webster, N.S. (2021). Integrating novel tools to elucidate the metabolic basis of microbial symbiosis in reef holobionts. Mar. Biol. 168, 175. 10.1007/s00227-021-03952-6.

93. Mos, B., and Dworjanyn, S.A. (2016). Early metamorphosis is costly and avoided by young, but physiologically competent, marine larvae. Mar. Ecol. Prog. Ser. 559, 117–129. 10.3354/meps11914.

94. Pechenik, J.A. (2006). Larval experience and latent effects-metamorphosis is not a new beginning. Integr. Comp. Biol. 46, 323–333. 10.1093/icb/icj028.

95. Edmunds, P.J., Cumbo, V.R., and Fan, T.-Y. (2013). Metabolic costs of larval settlement and metamorphosis in the coral *Seriatopora caliendrum* under ambient and elevated pCO2. J. Exp. Mar. Bio. Ecol. 443, 33–38. 10.1016/j.jembe.2013.02.032.

96. Grasso, L.C., Negri, A.P., Fôret, S., Saint, R., Hayward, D.C., Miller, D.J., and Ball, E.E. (2011). The biology of coral metamorphosis: molecular responses of larvae to inducers of settlement and metamorphosis. Dev. Biol. 353, 411–419. 10.1016/j.ydbio.2011.02.010.

97. Gilis, M., Meibom, A., Domart-Coulon, I., Grauby, O., Stolarski, J., and Baronnet, A. (2014). Biomineralization in newly settled recruits of the scleractinian coral *Pocillopora damicornis*. J. Morphol. 275, 1349–1365. 10.1002/jmor.20307.

98. Amjad, S., Nisar, S., Bhat, A.A., Shah, A.R., Frenneaux, M.P., Fakhro, K., Haris, M., Reddy, R., Patay, Z., Baur, J., et al. (2021). Role of NAD+ in regulating cellular and metabolic signaling pathways. Mol Metab 49, 101195. 10.1016/j.molmet.2021.101195.

99. Rivest, E.B., and Hofmann, G.E. (2015). Effects of temperature and pCO2 on lipid use and biological parameters of planulae of *Pocillopora damicornis*. J. Exp. Mar. Bio. Ecol. 473, 43–52. 10.1016/j.jembe.2015.07.015

100. Leuzinger, S., Willis, B.L., and Anthony, K.R.N. (2012). Energy allocation in a reef coral under varying resource availability. Mar. Biol. 159, 177–186. 10.1007/s00227-011-1797-1.

101. Neder, M., Saar, R., Malik, A., Antler, G., and Mass, T. (2022). New Insights on the Diurnal Mechanism of Calcification in the Stony Coral, Stylophora pistillata. Front. Mar. Sci. 8, 745171. 10.3389/fmars.2021.745171

102. Edmunds, P.J. (2012). Effect of pCO2 on the growth, respiration, and photophysiology of massive *Porites* spp. in Moorea, French Polynesia. Mar. Biol. 159, 2149–2160. 10.1007/s00227-012-2001-y

103. Richmond, R.H. (1987). Energetics, competency, and long-distance dispersal of planula larvae of the coral *Pocillopora damicornis*. Mar. Biol. 93, 527–533. 10.1007/BF00392790.

104. Nesa, B., Baird, A.H., Harii, S., Yakovleva, I., and Hidaka, M. (2012). Algal symbionts increase DNA damage in coral planulae exposed to sunlight. Zool. Stud. 51, 6.

105. Baird, A.H., Yakovleva, I.M., Harii, S., Sinniger, F., and Hidaka, M. (2021). Environmental constraints on the mode of symbiont transmission in corals. J. Exp. Mar. Bio. Ecol. 538, 151499. 10.1016/j.jembe.2020.151499.

106. Kitchen, S.A., Jiang, D., Harii, S., Satoh, N., Weis, V.M., and Shinzato, C. (2022). Coral larvae suppress heat stress response during the onset of symbiosis decreasing their odds of survival. Mol. Ecol. 31, 5813–5830. 10.1111/mec.16708.

107. Matthews, J.L., Crowder, C.M., Oakley, C.A., Lutz, A., Roessner, U., Meyer, E., Grossman, A.R., Weis, V.M., and Davy, S.K. (2017). Optimal nutrient exchange and immune responses operate in partner specificity in the cnidarian-dinoflagellate symbiosis. Proc. Natl. Acad. Sci. U. S. A. 114, 13194–13199. 10.1073/pnas.1710733114.

108. Kitchen, S.A., and Weis, V.M. (2017). The sphingosine rheostat is involved in the cnidarian heat stress response but not necessarily in bleaching. J. Exp. Biol. 220, 1709–1720. 10.1242/jeb.153858.

109. Ferrier-Pagès, C., Sauzéat, L., and Balter, V. (2018). Coral bleaching is linked to the capacity of the animal host to supply essential metals to the symbionts. Glob. Chang. Biol. 24, 3145–3157. 10.1111/gcb.14141

110. Hillyer, K.E., Dias, D.A., Lutz, A., Roessner, U., and Davy, S.K. (2017). Mapping carbon fate during bleaching in a model cnidarian symbiosis: the application of 13C metabolomics. New Phytol. 214, 1551–1562. 10.1111/nph.14515

111. Hillyer, K.E., Dias, D.A., Lutz, A., Wilkinson, S.P., Roessner, U., and Davy, S.K. (2017). Metabolite profiling of symbiont and host during thermal stress and bleaching in the coral *Acropora aspera*. Coral Reefs 36, 105–118. 10.1007/s00338-016-1508-y

112. Lee, H., Calvin, K., Dasgupta, D., Krinner, G., Mukherji, A., Thorne, P., Trisos, C., Romero, J., Aldunce, P., Barret, K., et al. (2023). IPCC, 2023: Climate Change 2023: Synthesis Report, Summary for Policymakers. Contribution of Working Groups I, II and III to the Sixth Assessment Report of the Intergovernmental Panel on Climate Change [Core Writing Team, H. Lee and J. Romero (eds.)]. IPCC, Geneva, Switzerland. (Intergovernmental Panel on Climate Change (IPCC)), p. 42.

113. Hartmann, A.C., Marhaver, K.L., Klueter, A., Lovci, M.T., Closek, C.J., Diaz, E., Chamberland, V.F., Archer, F.I., Deheyn, D.D., Vermeij, M.J.A., et al. (2019). Acquisition of obligate mutualist symbionts during the larval stage is not beneficial for a coral host. Mol. Ecol. 28, 141–155. 10.1111/mec.14967

114. Huffmyer, A.S. (2023). Cross-scale and multi-omic integration reveals transitions to symbiotic nutritional integration across coral ontogeny. Preprint at Open Science Framework, 10.17605/OSF.IO/KMU7H.

115. Padilla-Gamiño, J.L., and Gates, R.D. (2012). Spawning dynamics in the Hawaiian reef-building coral *Montipora capitata*. Mar. Ecol. Prog. Ser. 449, 145–160. 10.3354/meps09530.

116. Padilla-Gamiño, J.L., Hédouin, L., Waller, R.G., Smith, D., Truong, W., and Gates, R.D. (2014). Sedimentation and the reproductive biology of the Hawaiian reef-building coral *Montipora capitata*. Biol. Bull. 226, 8–18. 10.1086/bblv226n1p8

117. Hancock, J.R., Barrows, A.R., Roome, T.C., Huffmyer, A.S., Matsuda, S.B., Munk, N.J., Rahnke, S.A., Drury, C. (2021). Coral husbandry for ocean futures: leveraging abiotic factors to increase survivorship, growth, and resilience in juvenile *Montipora capitata*. Mar. Ecol. 657, 123–133. 10.3354/meps13534

118. Edmunds, P.J., and Davies, P.S. (1988). Post-illumination stimulation of respiration rate in the coral Porites porites. Coral Reefs 7, 7–9. 10.1007/BF00301975

119. Gibbin, E.M., Putnam, H.M., Gates, R.D., Nitschke, M.R., and Davy, S.K. (2015). Species-specific differences in thermal tolerance may define susceptibility to intracellular acidosis in reef corals. Mar. Biol. 162, 717–723. 10.1007/s00227-015-2617-9.

120. Lohman, B.K., Weber, J.N., and Bolnick, D.I. (2016). Evaluation of TagSeq, a reliable low-cost alternative for RNAseq. Mol. Ecol. Resour. 16, 1315–1321. 10.1111/1755-0998.12529.

121. Andrews, S. (2010). FastQC: A Quality Control Tool for High Throughput Sequence Data. http://www.bioinformatics.babraham.ac.uk/projects/fastqc/.

122. Ewels, P., Magnusson, M., Lundin, S., and Käller, M. (2016). MultiQC: summarize analysis results for multiple tools and samples in a single report. Bioinformatics 32, 3047–3048. 10.1093/bioinformatics/btw354

123. Chen, S., Zhou, Y., Chen, Y., and Gu, J. (2018). fastp: an ultra-fast all-in-one FASTQ preprocessor. Bioinformatics 34, i884–i890. 10.1093/bioinformatics/bty560

124. Stephens, T.G., Lee, J., Jeong, Y., Yoon, H.S., Putnam, H.M., Majerová, E., and Bhattacharya, D. (2022). High-quality genome assembles from key Hawaiian coral species. Gigascience 11, giac098. 10.1093/gigascience/giac098.

125. Kim, D., Paggi, J.M., Park, C., Bennett, C., and Salzberg, S.L. (2019). Graph-based genome alignment and genotyping with HISAT2 and HISAT-genotype. Nat. Biotechnol. 37, 907–915. 10.1038/s41587-019-0201-4

126. Kovaka, S., Zimin, A.V., Pertea, G.M., Razaghi, R., Salzberg, S.L., and Pertea, M. (2019). Transcriptome assembly from long-read RNA-seq alignments with StringTie2. Genome Biol. 20, 278. 10.1186/s13059-019-1910-1

127. Pertea, G., and Pertea, M. (2020). GFF Utilities: GffRead and GffCompare. F1000Res. *9*, 304. 10.12688/f1000research.23297.2

128. Williams, A., Chiles, E.N., Conetta, D., Pathmanathan, J.S., Cleves, P.A., Putnam, H.M., Su, X., and Bhattacharya, D. (2021). Metabolomic shifts associated with heat stress in coral holobionts. Sci Adv 7, eabd4210. 10.1126/sciadv.abd4210.

129. Su, X., Chiles, E., Maimouni, S., Wondisford, F.E., Zong, W.-X., and Song, C. (2020). In-Source CID Ramping and Covariant Ion Analysis of Hydrophilic Interaction Chromatography Metabolomics. Anal. Chem. 92, 4829–4837. 10.1021/acs.analchem.9b04181.

130. Melamud, E., Vastag, L., and Rabinowitz, J.D. (2010). Metabolomic analysis and visualization engine for LC-MS data. Anal. Chem. 82, 9818–9826. 10.1021/ac1021166

131. Lajeunesse, T.C., and Trench, R.K. (2000). Biogeography of two species of *Symbiodinium* (Freudenthal) inhabiting the intertidal sea anemone *Anthopleura elegantissima* (Brandt). Biol. Bull. 199, 126–134. 10.2307/1542872.

132. Camp, E.F., Suggett, D.J., Pogoreutz, C., Nitschke, M.R., Houlbreque, F., Hume, B.C.C., Gardner, S.G., Zampighi, M., Rodolfo-Metalpa, R., and Voolstra, C.R. (2020). Corals exhibit distinct patterns of microbial reorganisation to thrive in an extreme inshore environment. Coral Reefs 39, 701–716. 10.1007/s00338-019-01889-3.

133. Apprill, A., McNally, S., Parsons, R., and Weber, L. (2015). Minor revision to V4 region SSU rRNA 806R gene primer greatly increases detection of SAR11 bacterioplankton. Aquat. Microb. Ecol. 75, 129–137. 10.3354/ame01753.

134. Olito, C., White, C.R., Marshall, D.J., and Barneche, D.R. (2017). Estimating monotonic rates from biological data using local linear regression. J. Exp. Biol. 220, 759–764. 10.1242/jeb.148775.

135. Fox, J., and Weisberg, S. (2018). An R Companion to Applied Regression Second. (SAGE Publications).

136. R Core Team (2022). R: A language and environment for statistical computing (R Foundation for Statistical Computing). https://www.r-project.org/

137. Ogle, D.H., Doll, J.C., Wheeler, A.P., and Dinno, A. (2025). FSA: Simple Fisheries Stock Assessment Methods. R package version 0.9.6. 10.32614/CRAN.package.FSA

138. Rueden, C.T., Schindelin, J., Hiner, M.C., DeZonia, B.E., Walter, A.E., Arena, E.T., and Eliceiri, K.W. (2017). ImageJ2: ImageJ for the next generation of scientific image data. BMC Bioinformatics 18, 529. 10.1186/s12859-017-1934-z

139. Schindelin, J., Arganda-Carreras, I., Frise, E., Kaynig, V., Longair, M., Pietzsch, T., Preibisch, S., Rueden, C., Saalfeld, S., Schmid, B., et al. (2012). Fiji: An open-source platform for biological-image analysis. Nat. Methods 9, 676–682. 10.1038/nmeth.2019

140. Isomura, N., and Nishihira, M. (2001). Size variation of planulae and its effect on the lifetime of planulae in three pocilloporid corals. Coral Reefs 20, 309–315. 10.1007/s003380100180.

141. Lenth, R. (2024). emmeans: Estimated Marginal Means, aka Least-Squares Means. R package version 1.10.2. 10.32614/CRAN.package.emmeans

142. Okansen, J., and, et al. (2017). vegan: Community Ecology Package. R package version 2.6-6.1. 10.32614/CRAN.package.vegan

143. Love, M.I., Huber, W., and Anders, S. (2014). Moderated estimation of fold change and dispersion for RNA-seq data with DESeq2. Genome Biol. 15, 550. 10.1186/s13059-014-0550-8

144. Pantano, L. (2024). DEGreport: Report of DEG analysis. R package version 1.40.0. 10.18129/B9.bioc.DEGreport

145. Young, M.D., Wakefield, M.J., Smyth, G.K., and Oshlack, A. (2010). Gene ontology analysis for RNA-seq: accounting for selection bias. Genome Biol. 11, R14. 10.1186/gb-2010-11-2-r14.

146. Carlson, M. (2019). org.Ce.eg.db: Genome wide annotation for Worm. R package version 3.19.1. 10.18129/B9.bioc.org.Ce.eg.db

147. Sayols, S. (2020). rrvgo: a Bioconductor package to reduce and visualize Gene Ontology terms. microPubl. Biol. 10.17912/micropub.biology.000811

148. Rohart, F., Gautier, B., Singh, A., and Lê Cao, K.-A. (2017). mixOmics: An R package for ’omics feature selection and multiple data integration. PLoS Comput. Biol. 13, e1005752. 10.1371/journal.pcbi.1005752.

149. Hervé, M. (2020). RVAideMemoire: Testing and Plotting Procedures for Biostatistics. R package version 0.9-83-7. 10.18129/B9.bioc.rrvgo

150. Galindo-Prieto, B., Eriksson, L., and Trygg, J. (2014). Variable influence on projection (VIP) for orthogonal projections to latent structures (OPLS). J. Chemom. 28, 623–632. 10.1002/cem.2627.

151. Pang, Z., Chong, J., Zhou, G., de Lima Morais, D.A., Chang, L., Barrette, M., Gauthier, C., Jacques, P.-É., Li, S., and Xia, J. (2021). MetaboAnalyst 5.0: narrowing the gap between raw spectra and functional insights. Nucleic Acids Res. 49, W388–W396. 10.1093/nar/gkab382.

152. McMurdie, P.J., and Holmes, S. (2013). phyloseq: an R package for reproducible interactive analysis and graphics of microbiome census data. PLoS One 8, e61217. 10.1371/journal.pone.0061217.

153. Schloss, P.D., Westcott, S.L., Ryabin, T., Hall, J.R., Hartmann, M., Hollister, E.B., Lesniewski, R.A., Oakley, B.B., Parks, D.H., Robinson, C.J., et al. (2009). Introducing mothur: open-source, platform-independent, community-supported software for describing and comparing microbial communities. Appl. Environ. Microbiol. 75, 7537–7541.

154. Quast, C., Pruesse, E., Yilmaz, P., Gerken, J., Schweer, T., Yarza, P., Peplies, J., and Glöckner, F.O. (2013). The SILVA ribosomal RNA gene database project: improved data processing and web-based tools. Nucleic Acids Res. 41, D590–D596. 10.1093/nar/gks1219.

155. Rognes, T., Fluori, T., Nichols, B., Quince, C., and Mahé, F. (2016). VSEARCH: a versatile open source tool for metagenomics. PeerJ, e2584. 10.7717/peerj.2584

