## Supplemental Figures for "Metabolic dynamics of the coral-algal symbiosis from fertilization to settlement identify critical coral energetic vulnerabilities"

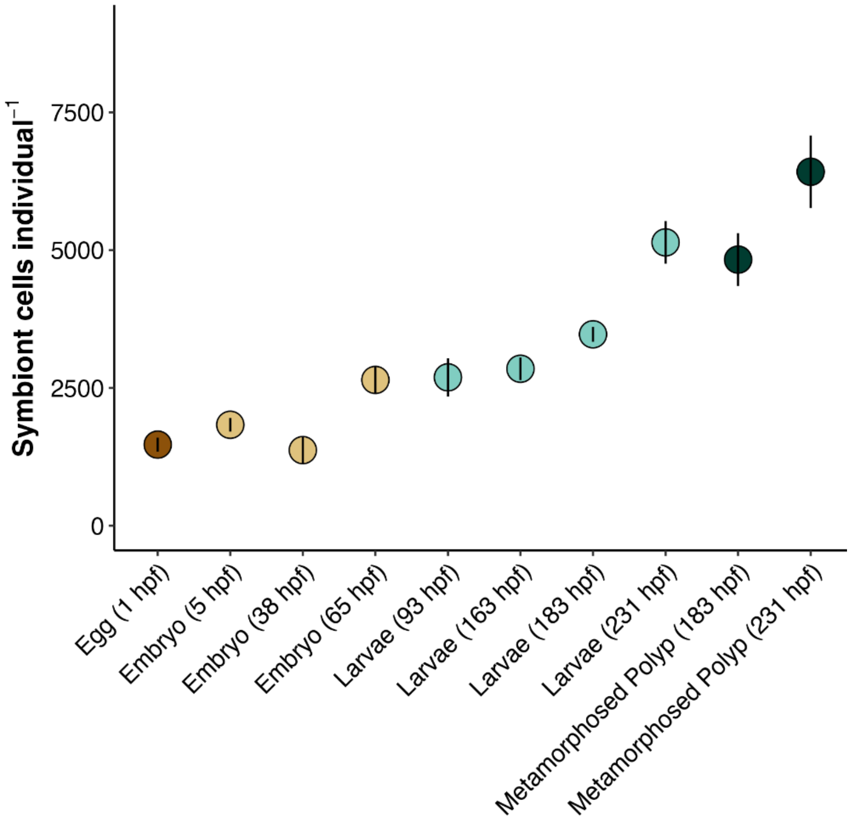


**Fig S1. Symbiont cell density per individual from egg to metamorphosed polyp stages.** Data are represented as mean ± standard error. Life stage is indicated on the x-axis as hours post fertilization (hpf). Color corresponds to major life history grouping (eggs=brown, embryos=yellow, larvae=cyan, metamorphosed polyps=green, attached recruits=pink).

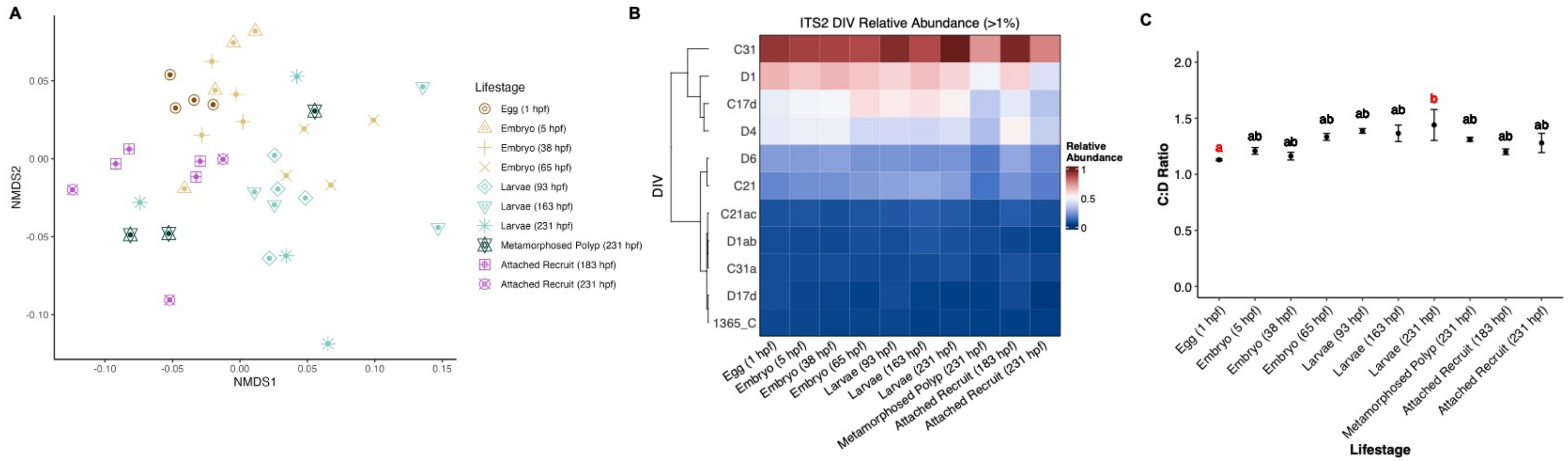


**Fig S2. Symbiodiniaceae ITS2 analysis.** (A) NMDS plot of Symbiodiniaceae communities across development (eggs=brown, embryos=yellow, larvae=cyan, metamorphosed polyps=green, attached recruits=pink). Shape indicates life stage. (B) Heatmap of ITS2 DIV relative abundance of taxa abundant at >1% relative abundance across life stages. Red = relative abundance of 1; Blue = relative abundance of 0. (C) Ratio of *Cladocopium* sp. to *Durusdinium glynii* relative abundance across life stages. Error bars indicate standard error of mean. Shared letters indicate groups are not significantly different. Groups with red letters are significantly different at post hoc $p<0.05$.

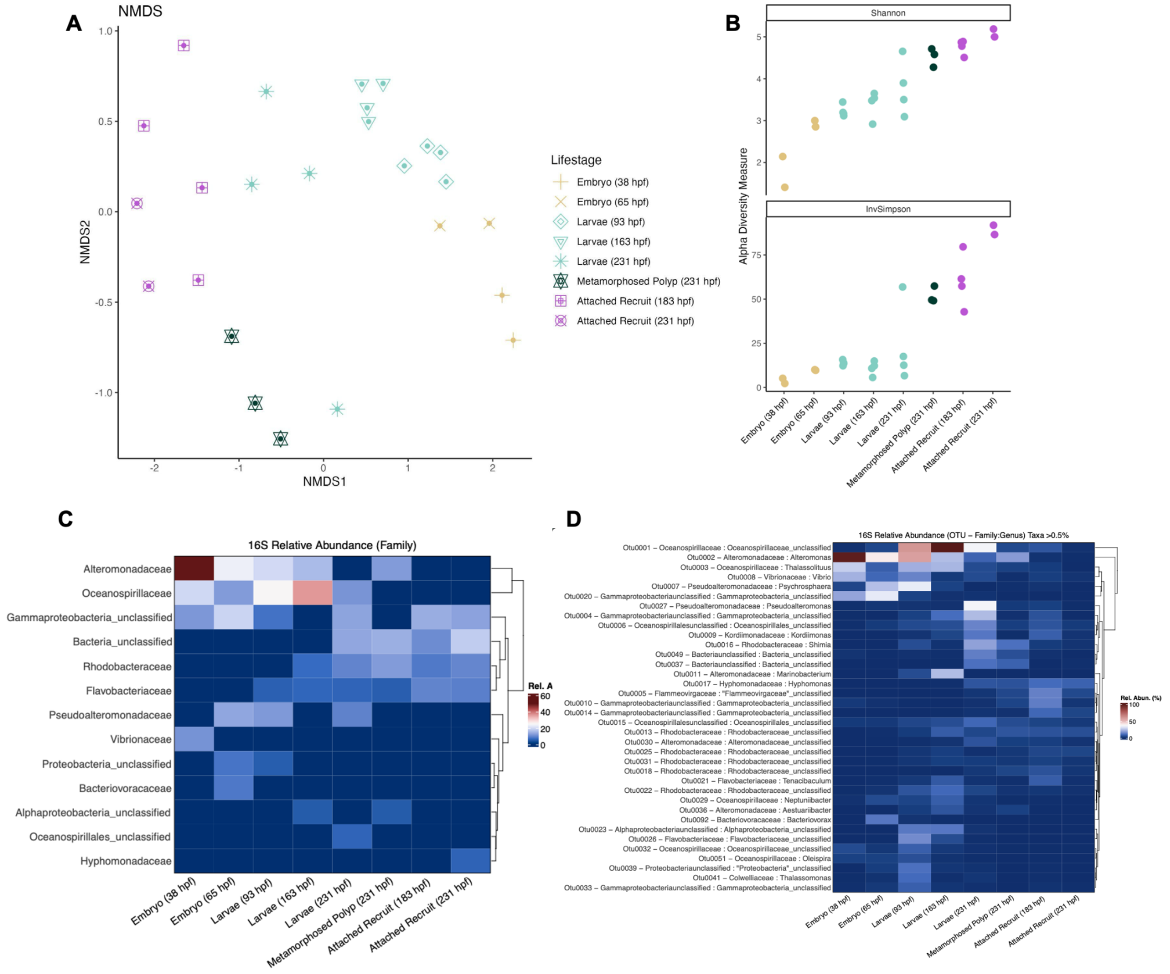


**Fig S3. Bacterial community 16S analysis.** (A) NMDS plot of bacterial communities across development (eggs=brown, embryos=yellow, larvae=cyan, metamorphosed polyps=green, attached recruits=pink). Shape indicates life stage. (B) Alpha diversity metrics of bacterial communities calculated as Shannon (top) and Inverse Simpson (bottom) metrics. (C) Heatmap of bacterial relative abundance across life stages at the phylum level. (D) Heatmap of bacterial relative abundance across life stages at the family:genus level in taxa at greater than 0.5% relative abundance. Red colors indicate higher relative abundance.

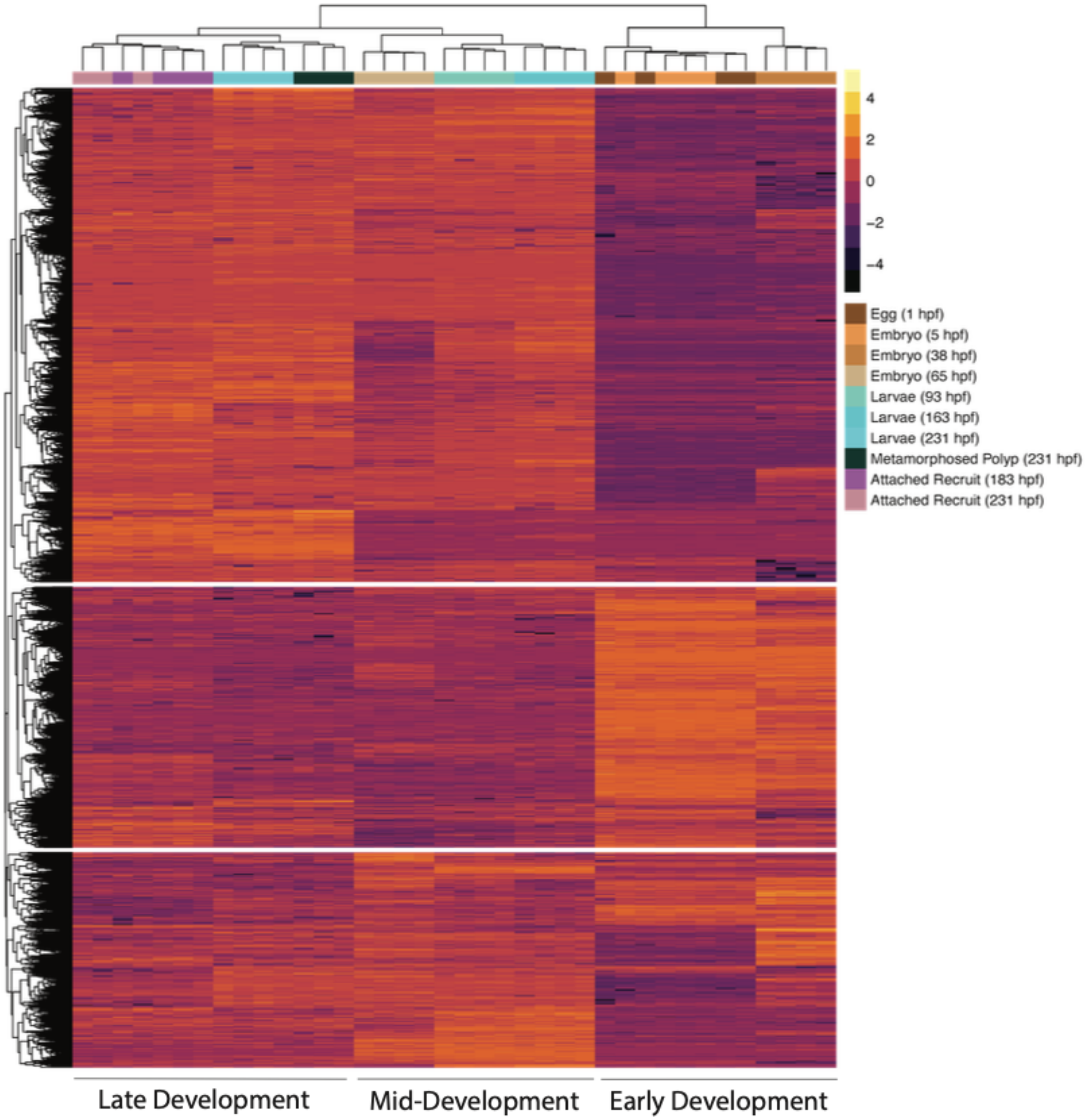


**Fig S4. Differentially expressed genes across development.** Heatmap of expression (z-score) of differentially expressed genes (9,181 total DEGs; y-axis) across time points (x-axis) identified by likelihood ratio tests (FDR P-value < 0.05). Within the heatmap, orange indicates increased expression with purple indicating reduced expression relative to the average of that gene's expression in all samples. Column annotation colors indicate life stages.

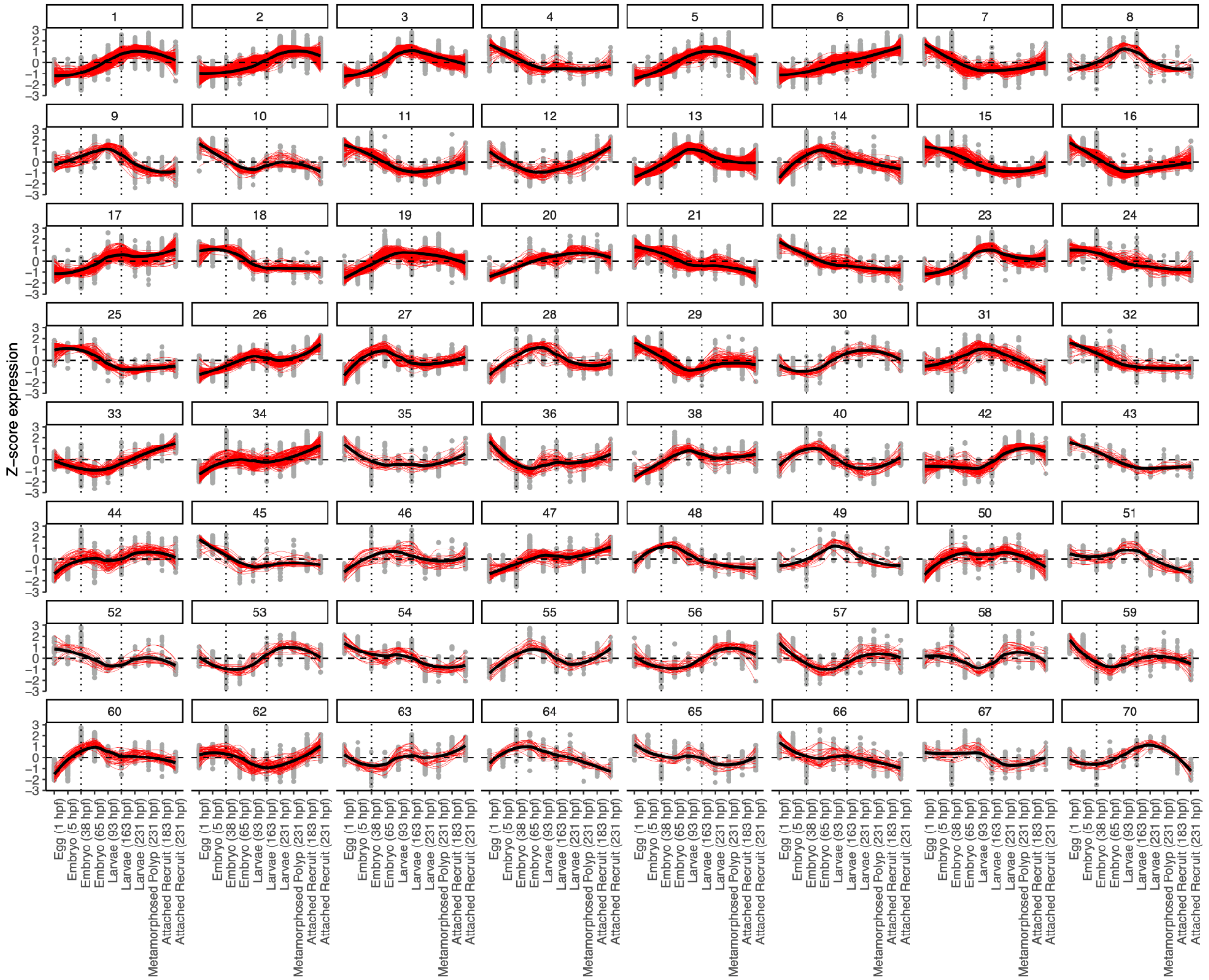


**Fig S5. Expression patterns of gene clusters across development.** Panels show detected clusters (minimum number of genes = 15) of genes that share expression patterns across life stages. Expression displayed as z-score of variance stabilized transformed gene counts. Dotted lines indicate 38 hpf and 163 hpf that separate early, mid, and late developmental stages. Clusters were assigned to developmental patterns based on the time at which expression peaked: early development (1-38 hpf), mid development (65-163 hpf) and late development (183-231 hpf). Red lines indicate expression for individual genes with black lines indicating the cluster mean expression across development.

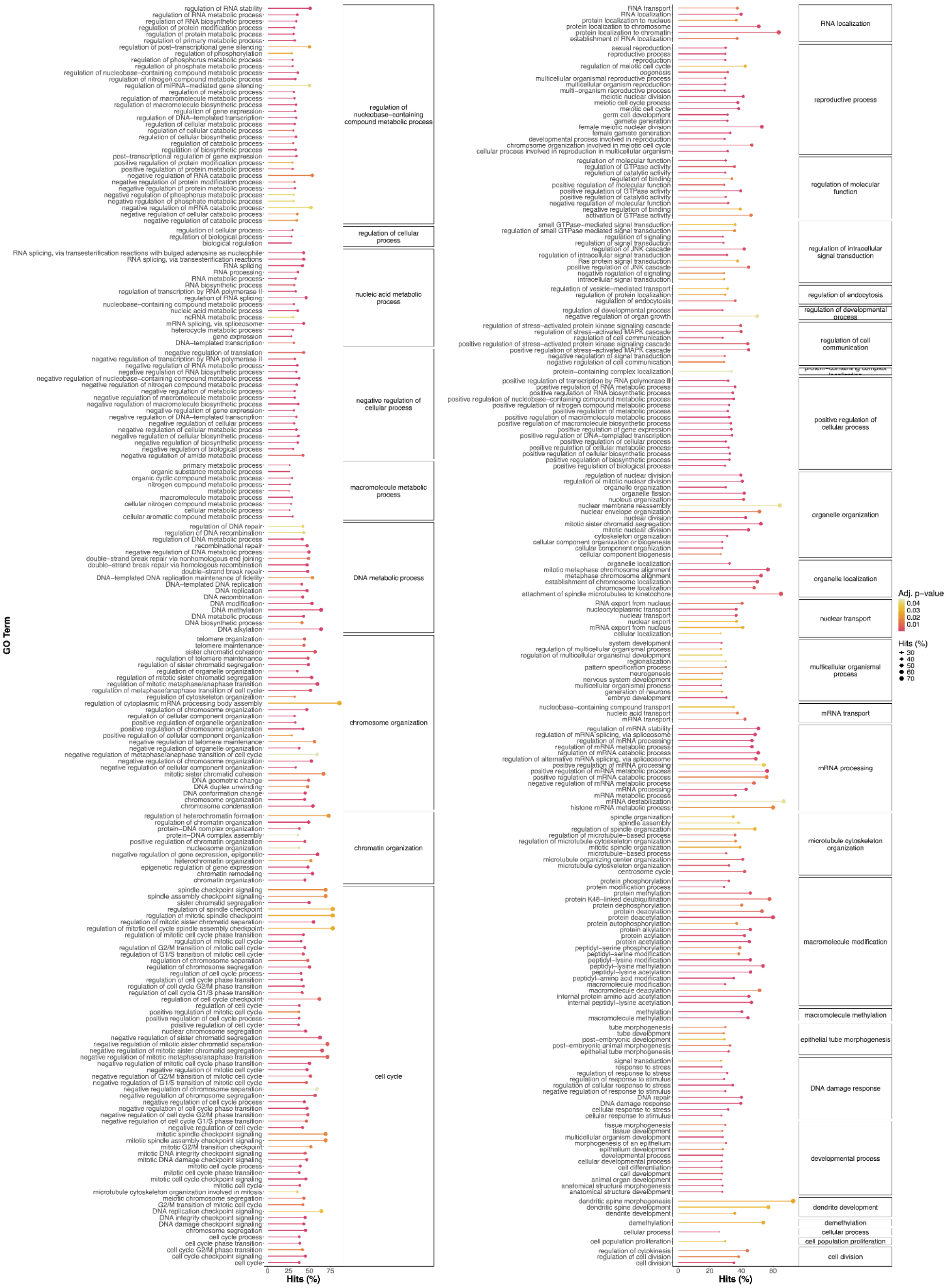


**Fig S6. Functional enrichment of genes in early development.** Lollipop plot of functional enrichment of genes peaking in early development (1-38 hpf). Plot shows significant (FDR P<0.05) gene ontology (GO) terms (Biological Process; BP). x-axis and dot size indicate % hits (percentage of the number of DEGs in the respective GO category compared to total number of genes included in that category) with color indicating P-value. Parent term BP is indicated in panels on the right and individual GO terms on the left.

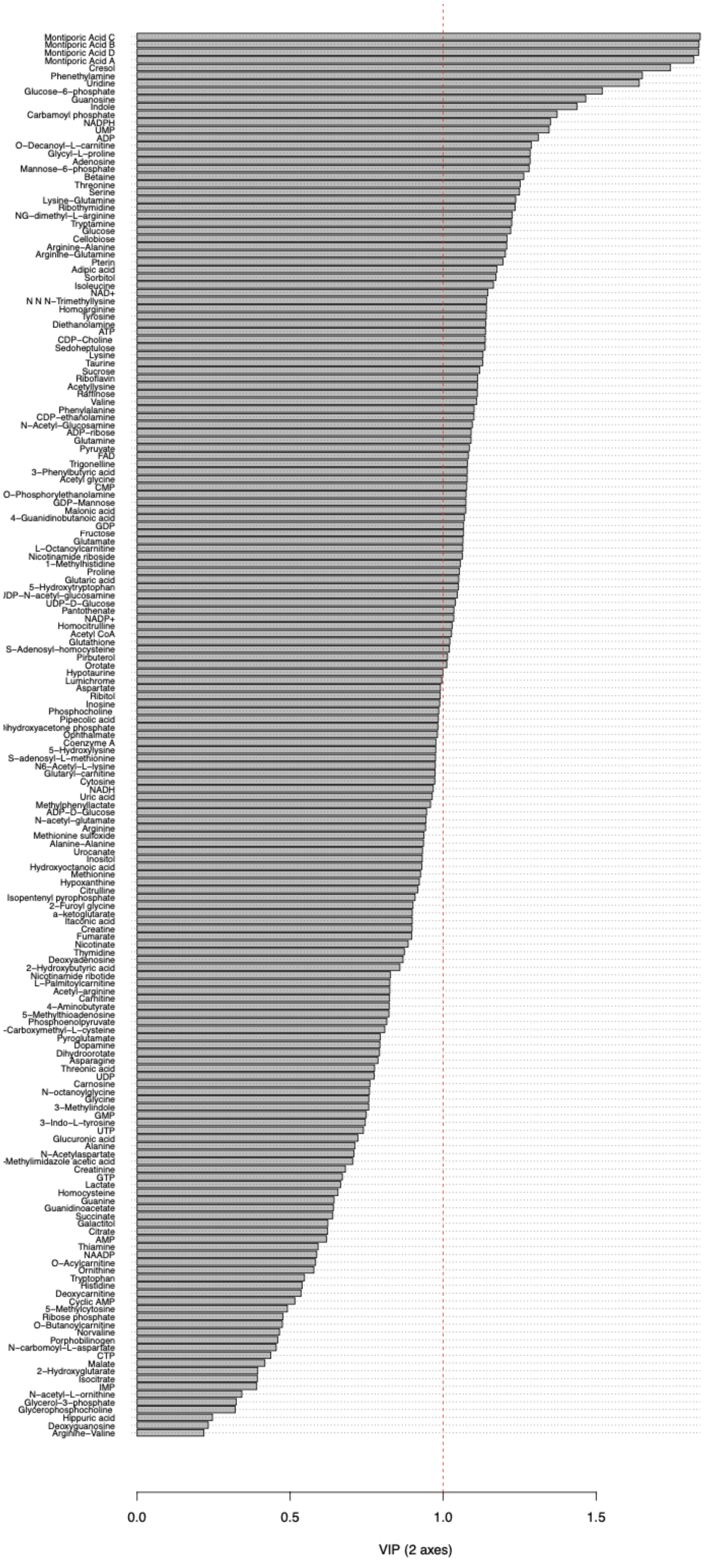


**Fig S7. Metabolite variables of importance in project (VIP) scores for all metabolites.** Red dashed line indicates VIP = 1, above which metabolites are statistically important in discriminating between life stage groups. VIP scores calculated for components 1 and 2 in the partial least squares discriminant model.
